# Enriching for orthologs increases support for Xenacoelomorpha and Ambulacraria sister relationship

**DOI:** 10.1101/2021.12.13.472462

**Authors:** Peter O Mulhair, Charley GP McCarthy, Karen Siu-Ting, Christopher J Creevey, Mary J O’Connell

**Author notes:** Both authors contributed equally to this work (Peter O Mulhair, Charley GP McCarthy).

## Abstract

Conflicting studies place a group of bilaterian invertebrates containing xenoturbellids and acoelomorphs, the Xenacoelomorpha, as either the primary emerging bilaterian phylum, or within Deuterostomia, sister to Ambulacraria. While their placement as sister to the rest of Bilateria supports relatively simple morphology in the ancestral bilaterian, their alternative placement within Deuterostomia suggests a morphologically complex ancestral Bilaterian along with extensive loss of major phenotypic traits in the Xenacoelomorpha. More recently, further studies have brought into question whether Deuterostomia should be considered monophyletic at all. Hidden paralogy presents a major challenge for reconstructing species phylogenies. Here we assess whether hidden paralogy has contributed to the conflict over the placement of Xenacoelomorpha. Our approach assesses previously published datasets, enriching for orthogroups whose gene trees support well resolved clans elsewhere in the animal tree of life. We find that the majority of constituent genes in previously published datasets violate incontestable clans, suggesting that hidden paralogy is rife at this depth. We demonstrate that enrichment for genes with orthologous signal alters the final topology that is inferred, whilst simultaneously improving fit of the model to the data. We discover increased, but ultimately not conclusive, support for the existence of Xenambulacraria in our orthology enriched set of genes. At a time when we are steadily progressing towards sequencing all of life on the planet, we argue that long-standing contentious issues in the tree of life will be resolved using smaller amounts of better quality data that can be modelled adequately.

## Introduction

Many studies have attempted to address the placement of Xenacoelomorpha using a mixture of molecular and morphological characters (Hejnol & Pang 2016; Haszprunar 2016; Ruiz-Trillo & Paps 2016). Traditionally, this group of small marine worms have been placed as the primary emerging bilaterian lineage (Figure 1A), implying a less complex body plan of the most recent common ancestor of Bilateria, with a simple brain, blind gut, and lacking excretory and vascular systems (Ruiz-Trillo et al. 2004; Paps et al. 2009; Ruiz-Trillo & Paps 2016; Hejnol & Pang 2016; Rouse et al. 2016; Cannon et al. 2016). The alternative placements within Deuterostomia (Figure 1B and C) suggest a secondary simplification of a large number of significant morphological traits in the Xenacoelomorph clade which are considered to be present in the last common ancestor of deuterostomes, i.e. the loss such as complex organ systems, a digestive system with mouth and anus, coeloms, and body compartmentalisation (Bourlat et al. 2003, 2006; Philippe, Brinkmann, Copley, et al. 2011; Philippe et al. 2019; Kapli & Telford 2020). More recently, the placement of Xenacoelomorpha within Bilateria has brought into question whether Deuterostomia should be considered monophyletic at all (Philippe et al. 2019; Kapli et al. 2021). The paraphyletic Deuterostomia hypothesis places Chordata as sister to Protostomia to the exclusion of Xenambulacraria (Xenacoelomorpha plus Ambulacraria) (Figure 1C), suggesting that the common ancestor of all Bilateria possessed many deuterostome traits such as radial cleavage and the development of the anus from the blastopore, and that these traits were significantly altered or lost at the emergence of Protostomia (Kapli et al. 2021). Empirical analyses of newly generated genomic and transcriptomic data continue to produce conflicting placements for Xenacoelomorpha, with support garnered for both the Nephrozoa (Figure 1A) (Cannon et al. 2016; Rouse et al. 2016) and the Xenambulacraria (Figure 1B & C) (Philippe et al. 2019) hypotheses.

**Figure 1:**
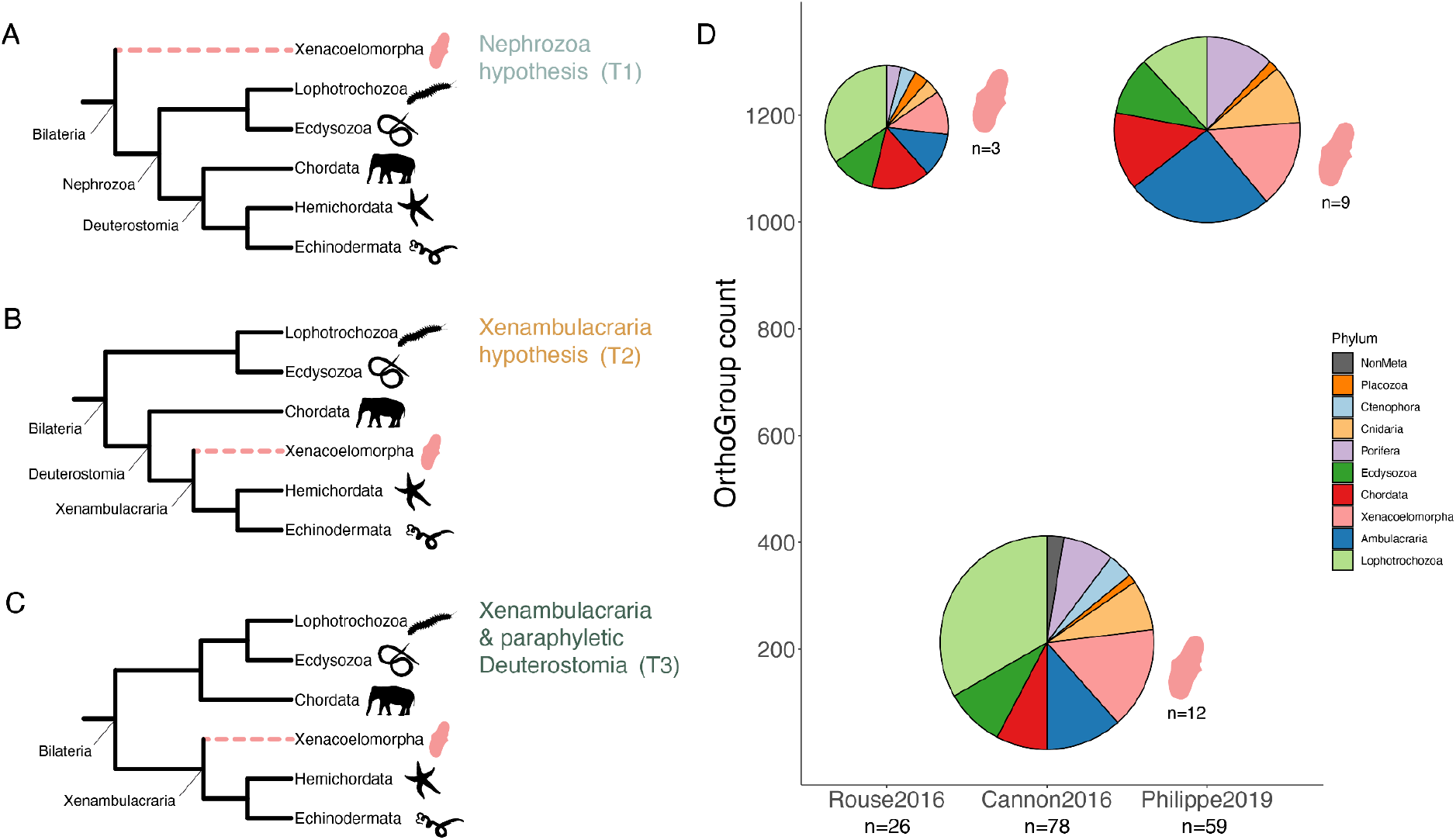
Conflicting placements for Xenacoelomorpha and distribution of phylum sampling in three major datasets applied to resolve their position. **(A)** Nephrozoa hypothesis (T1) posits that Xenacoelomorpha are sister to the remaining bilaterian phyla. **(B)** Xenambulacraria hypothesis (T2) places Xenacoelomorpha as sister to Ambulacraria, a clade consisting of Hemichordata and Echinodermata. **(C)** Xenambulacraria has also been suggested along with non-monophyletic Deuterostomia, with Chordata placed sister to Protostomia. **(D)** Overview of previous phylogenomic studies addressing the placement of Xenacoelomorpha. Three studies are shown, with circles sized based on the number of taxa sampled for the analysis. Each of the wedges in the circle are coloured based on a given animal phylum (legend) and sized based on the number of species in that phylum that were used in the study. The y-axis represents the number of orthogroups used in the phylogenomic analysis.

Finding the definitive placement of Xenacoelomorpha is complicated by biological and systematic features of the phylum including high rates of sequence evolution and gene loss within Xenacoelomorpha (particularly within Acoelomorpha), and short internal branches in the deep splitting lineages of Bilateria (Philippe et al. 2019; Kapli et al. 2021). Careful analyses of novel and previously published datasets (Cannon et al. 2016; Rouse et al. 2016) by Philippe et al (2019) attempted to address some of these features by using site heterogeneous models, data re-coding, and the ‘best’ gene sets based on signal within the gene trees. In doing so they recovered Xenacoelomorpha sister to Ambulacraria plus a paraphyletic Deuterostomia (T3, Figure 1C). Additionally, Kapli and Telford (2020) analysed the effects of model misspecification on the placement of Xenacoelomorpha and found that misfitting site homogeneous models result in the Nephrozoa hypothesis (T1, Figure 1A) as a result of systematic bias caused by long branch attraction (Kapli & Telford 2020) rather than true evolutionary signal. However, these analyses, and all previous studies addressing this phylogenetic question, did not explore the distribution and strength of phylogenetic signal in the underlying data. To date, there have been no assessments of misleading signal caused by hidden paralogy, and whether accounting for this increases support for the placement of Xenacoelomorpha in the animal tree of life.

Assessing how various biases in data such as rate and compositional heterogeneity may affect resulting topology has seen model adequacy and fit receive a justifiable level of attention (Morgan et al. 2013; Feuda et al. 2017; Philippe et al. 2019; Kapli & Telford 2020; Redmond & McLysaght 2021). However, the underlying signal within large datasets for phylogeny reconstruction warrants further exploration. Most of these large scale studies set out to reduce paralogy, yet a standardised assessment of the prevailing levels of hidden paralogy is lacking. Hidden paralogy (Doolittle & Brown 1994) is driven by gene duplication followed by subsequent differential loss, and has been shown to have profound effects on species tree inference (Supplementary Figure S1A) (Brown & Thomson 2017; Siu-Ting et al. 2019; Walker et al. 2020; Natsidis et al. 2021). Given the observed high rates of genome duplication and gene turnover in animals (Grau-Bové et al. 2017; Richter et al. 2018; Paps & Holland 2018; Fernández & Gabaldón 2020; Guijarro-Clarke et al. 2020), combined with the use of incomplete transcriptomic and genomic data in phylogenomic analyses, hidden paralogy and misleading phylogenetic signal remains a significant concern for species tree inference.

Here, we consider three previously published datasets which supported two of the three placements of Xenacoelomorpha (Cannon et al. 2016; Rouse et al. 2016; Philippe et al. 2019). We filter gene families based on their ability to resolve well known clans (excluding contentious nodes of interest) (Siu-Ting et al. 2019) and measure the distribution of support for the alternative placements of Xenacoelomorpha between the genes that passed and failed this filter. We assess whether filtering these datasets for orthologs in this way reduces conflict, improves model fit, and increases support for a single topology. Finally we demonstrate that applying this additional filter can result in significant changes to the species topology recovered.

## Results

### Large variation between dataset sizes and taxon sampling

We set out to analyse the distribution of phylogenetic signal within each of the three datasets from Rouse et al. (2016), Cannon et al. (2016), and Philippe et al. (2019), these datasets are referred to as Rouse2016, Cannon2016 and Philippe2019 throughout. The datasets differ in their taxon sampling, data filtering, phylogenomic methods applied, and ultimately, in their placement of Xenacoelomorpha (Figure 1D, Table 1). The sample size differs significantly between studies, a factor which is known to affect phylogenetic inference (Zwickl & Hillis 2002; Wilberg 2015; Philippe, Brinkmann, Lavrov, et al. 2011), with 26 species (including 3 Xenacoelomorpha species), 78 species (12 Xenacoelomorpha), and 59 species (9 Xenacoelomorpha) sampled in Rouse2016, Cannon2016, and Philippe2019 respectively (Figure 1D). The phylogenetic spread of taxa sampling also differs across the three studies, with the most even spread across animal phyla observed in Philippe2019 (Figure 1D). The underlying sources of sequence data used in each of these studies differ, and the ratio of transcriptomes to genomes used vary significantly between studies (Table 1). It is known that transcriptomic data, or indeed poor quality genomic data, may negatively affect phylogenetic inference (Brown & Thomson 2017; Siu-Ting et al. 2019; Spillane et al. 2021). Additionally, the data matrices used in these three studies were constructed using different orthology assignment tools (i.e. Agalma (Dunn et al. 2013), HaMStR (Ebersberger et al. 2009) and OMA (Altenhoff et al. 2019)). Therefore, the resulting data matrices consist of different numbers of genes and sites (Table 1). Methods for data filtering also vary, with Cannon et al. (2016) and Philippe et al. (2019) applying different approaches to reduce paralogs and Cannon et al. (2016) explicitly filtering for saturation and compositional heterogeneity (although Philippe et al, (2019) do employ data re-coding as a *post-hoc* solution).

**Table 1:**
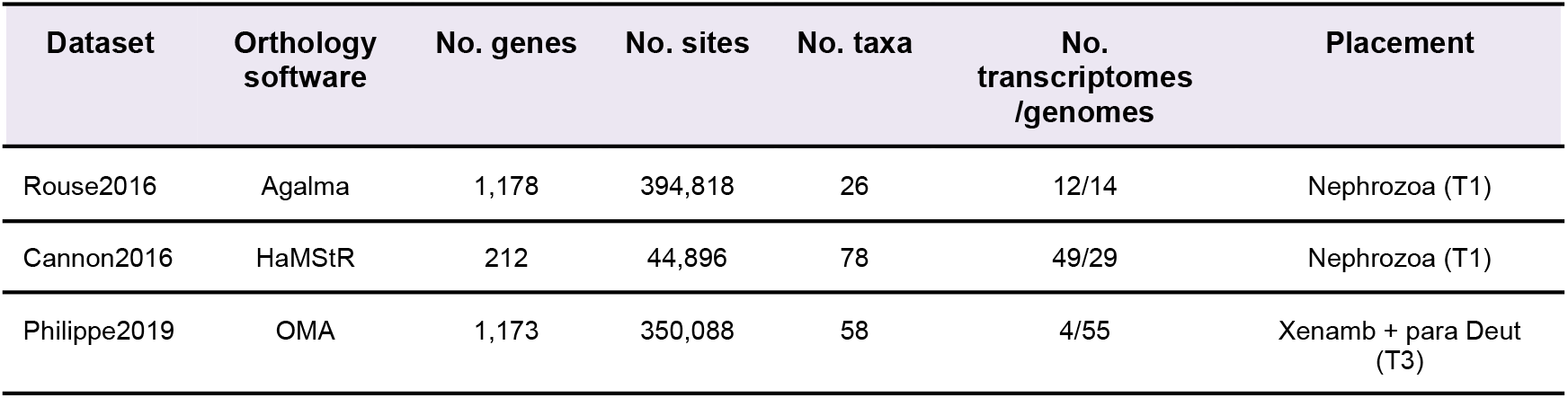
Overview of previously published datasets used in current analysis. The three datasets Rouse2016 (Rouse et al. 2016), Cannon2016 (Cannon et al. 2016) and Philippe2019 (Philippe et al. 2019) included in this study are detailed. The Orthology detection software used is detailing, along with the number of genes (No. genes), the number of sites in the alignment (No. sites), the number of taxa (No. taxa) and the proportion of the data that was from transcriptome or genome sequencing (No. transcriptomes/genomes). The placement resolved in the original publication is detailed in the final column (Placement). Xenamb + para Deut = Xenambulacraria and Paraphyletic Deuterostomes.

| Dataset | Orthology software | No. genes | No. sites | No. taxa | No. transcriptomes /genomes | Placement |
| --- | --- | --- | --- | --- | --- | --- |
| Rouse2016 | Agalma | 1,178 | 394,818 | 26 | 12/14 | Nephrozoa (T1) |
| Cannon2016 | HaMStR | 212 | 44,896 | 78 | 49/29 | Nephrozoa (T1) |
| Philippe2019 | OMA | 1,173 | 350,088 | 58 | 4/55 | Xenamb + para Deut (T3) |

### Frequent violation of known monophyletic clans across previously published datasets suggests inadvertent paralog inclusion

For each of the three datasets (Philippe2019, Rousse2016 and Canon2016) we used Clan_Check (Siu-Ting et al. 2019) to assess whether the constructed gene trees could recapitulate known, incontestable clans (*sensu* (Wilkinson et al. 2007)) (Supplementary Figure S1). Under the assumption that a tree inferred from a set of orthologous genes should recapitulate incontestable clans (Supplementary Figure S2), trees that consistently violate these assumptions with high support may indicate inadvertent paralog selection. The incontestable clans we defined are Porifera, Ctenophora, Cnidaria, Bilateria, Protostomia, Deuterostomia, Xenacoelomorpha, Ambulacraria, Lophotrochozoa, Ecdysozoa, and Chordata. We found widespread violation of the tested clans in gene trees from each of the three datasets (Figure 2A). For example, in the Philippe et al. (2019) dataset, each clan was violated by between 60-99% of the gene trees (Figure 2A). The clan which was violated most often was Deuterostomia, with 1,155/1,164 (99%) of the gene trees failing to recapitulate the clan. This is of particular interest due to recent questioning of the true monophyly of this major group (Philippe et al. 2019, Kapli and Telford 2021). Similar patterns are observed in each of the other two datasets (*i.e*. Rouse2016 and Canon2016), with widespread violation of incontestable clans in the constructed gene trees (Figure 2A). We classified gene families as those with extensive violation of clans by Clan_check (“CC fail”) and those with low or no violations (“CC pass”). For the CC pass gene set we considered only those gene families that could recapitulate at least 4, 3, and 5 clans for Rouse2016, Cannon2016, and Philippe2019, respectively (Supplementary Table S1)(see materials and methods). This resulted in three datasets of 70, 16, and 65 genes for Rouse2016, Cannon2016 and Philippe2019, respectively (Figure 2B).

**Figure 2:**
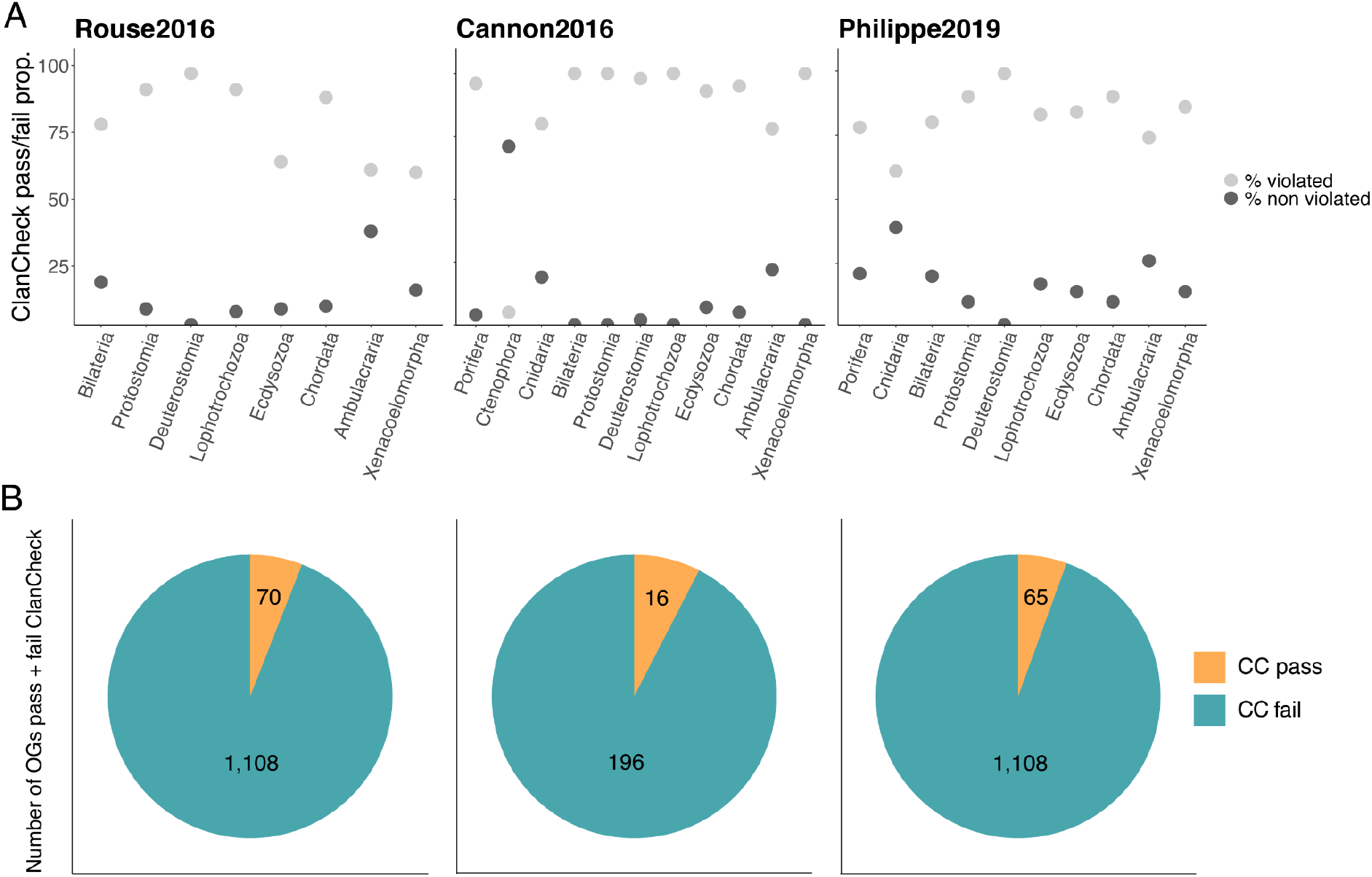
Results of Clan_Check Filtering for Rouse2016, Cannon2016 and Philippe2019 datasets. **(A)** The proportion of gene trees violating each of the clans tested for each of the three datasets **(B)** The number of genes from the original dataset which passed or failed the Clan_Check filter. “CC pass” (yellow) genes are those gene families that could recapitulate above the threshold number of the splits and “CC fail” (blue) are those that failed to recapitulate enough clans above the threshold.

### Clan_Check filtering improves overall phylogenetic signal without biasing for compositional heterogeneity, rates of sequence evolution, or function

We compared overall branch lengths, compositional heterogeneity, and gene function between the CC fail and CC pass gene sets for each of the three datasets (Rouse2016, Cannon2016 and Philippe2019). We found no significant difference between the two sets of genes for any of the traits tested in any of the three datasets (Supplementary Table S2). This suggests that our filter does not generate gene sets with biases in branch length, compositional heterogeneity, or function (Siu-Ting et al. 2019). Additionally, we also measured taxon sampling between the genes that failed and passed the Clan_Check filter and found no evidence for subsampling of genes with biases for species in certain phyla (Supplementary Figure S3).

Additionally, we used seven different gene and tree based metrics (see materials and methods) shown to reflect the accuracy of species tree inference, to assess whether the dataset enriched for orthologs, *i.e*. CC pass, differ significantly from the dataset that failed the Clan_Check test, *i.e*. CC fail (Shen et al. 2016, 2021). We found that for all three datasets, and across almost all tests, the smaller dataset of genes enriched for orthologs (CC pass) contained a significantly greater amount of phylogenetic signal based on the different metrics (Wilcoxon rank-sum test; p<0.05) (Figure 3). Some deviations were noted in the “long branch score”, “saturation” and “treeness divided by relative composition variability” in some datasets. Overall, however, when we filter each of the three datasets based on their ability to recapitulate known clans, phylogenetic signal is increased, providing justification for reducing the data matrices for phylogenetic inference.

**Figure 3:**
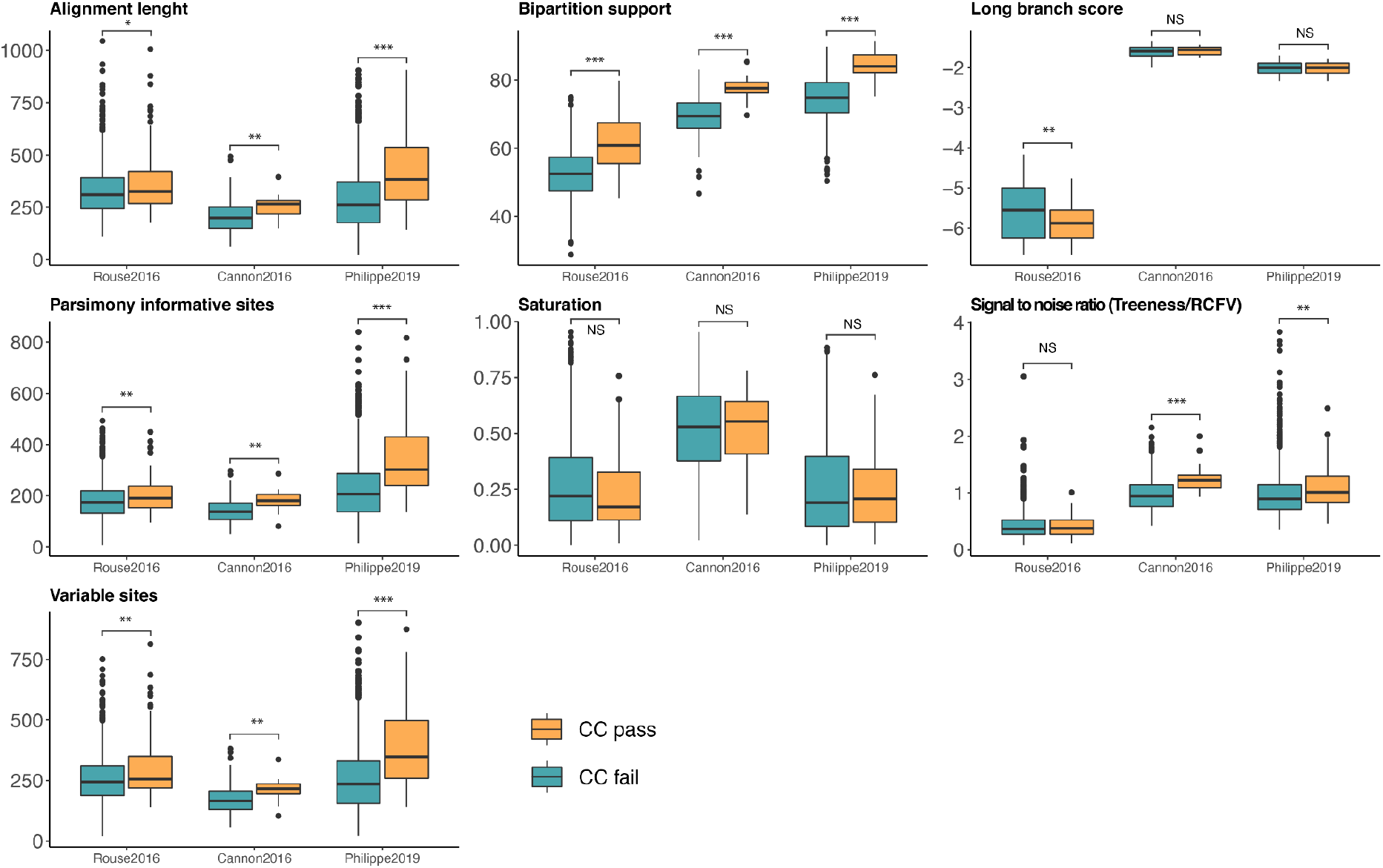
Summary of seven tree based traits of signal between CC fail and pass OGs for each of the three datasets. Metrics tested included alignment length, bipartition support, long branch score, number of parsimony informative sites, level or saturation, treeness divided by relative compositional variability (RCFV), and the number of variable sites. Wilcoxon rank-sum test was then carried out between CC pass and CC fail gene sets for each metric tested for each of the three datasets (NS = *P*-value > 0.05, \**P*-value < 0.05, \*\**P*-value < 0.01, \*\*\**P*-value < 0.001).

### Gene level signal varies between datasets and support for a single topology increases in filtered datasets

We determined the proportion of support for each of the three currently debated positions for the Xenacoelomorpha; T1, T2, and T3 (Figure 1A-C) from the set of CC-pass and CC-fail genes from Rouse2016, Cannon2016 and Philippe2019. Gene-wise log likelihood values for the full set of genes in each dataset showed T1, Nephrozoa hypothesis to be the dominant hypothesis in Rouse2016 (47% of genes) and Cannon2016 (58% of genes), but found significantly reduced support for T1 in Philippe2019 (28% genes) (Figure 4). This shows clear distinction in signal between the datasets, which also reflect the hypothesis supported in the original studies. It is worth noting that the signal for both Xenambulacraria hypotheses (i.e. T2 and T3) represents a majority in Rouse2016 (53% of genes) in comparison to T1. Comparing the gene-wise signal between the full gene sets and the set of CC-pass genes, we find increased support for T1 (Nephrozoa hypothesis) in both Rouse2016 (57% of genes) and Cannon2016 (69% of genes) in the CC-pass gene set (Figure 4). In contrast, we find a further decrease in support for T1 in the CC-pass gene set of Philippe2019 (12% of genes) and an increase of support for both T2 (43% of genes) and T3 (45% of genes).

**Figure 4:**
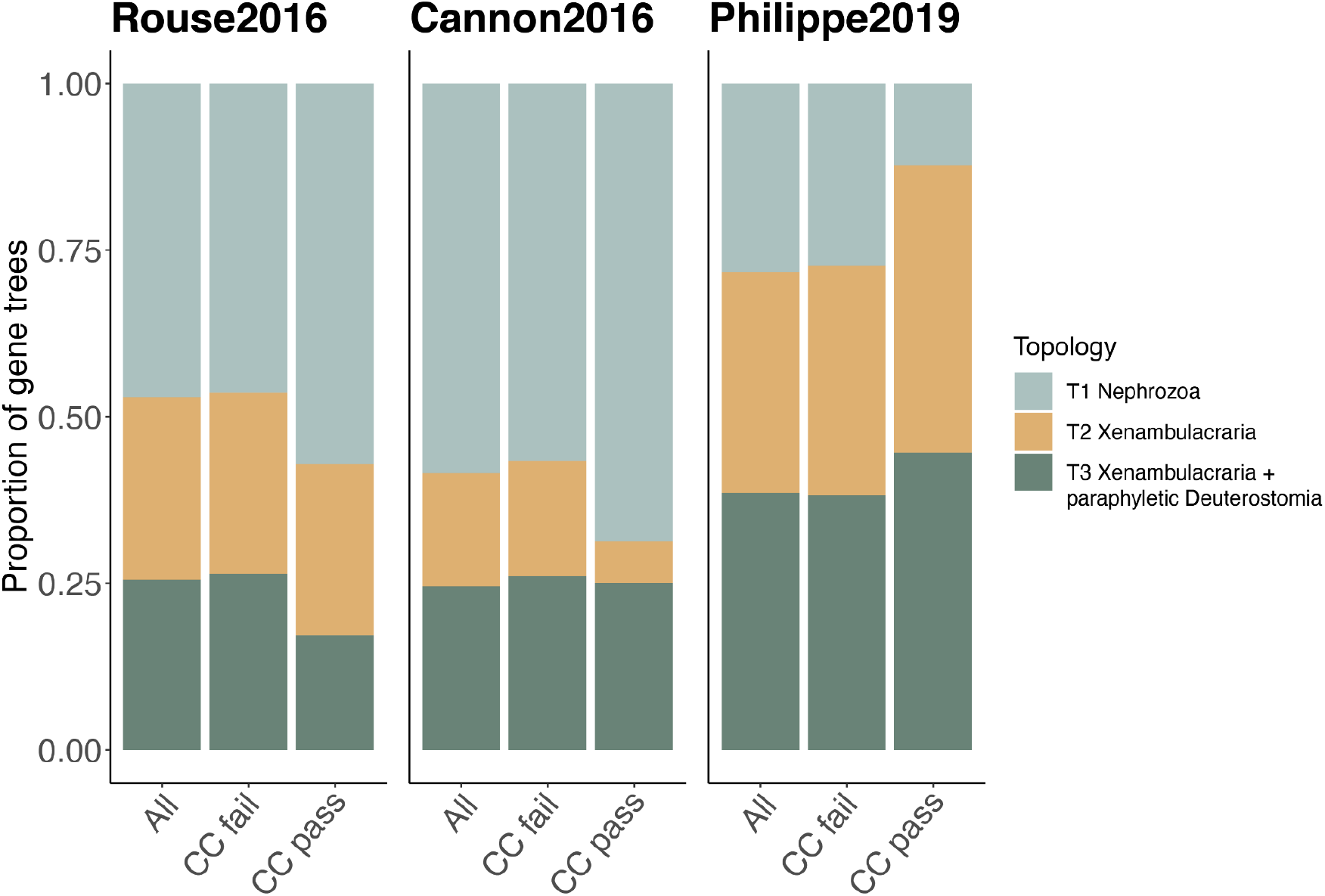
Gene-wise support for alternative topologies from gene trees that pass or fail the Clan_Check filter for each of the three datasets. The x-axis represents all genes along with the CC fail or CC pass subsets of the data for each of the three datasets, Rouse2016, Cannon2016 and Philippe2019 (left to right). The y-axis represents the proportion of the data that supports each of the three conflicting positions for Xenacoelomorpha: T1 = Nephrozoa hypothesis (yellow), T2 = Xenambulacraria (red), and T3 = Xenambulacraria with paraphyletic Deuterostomes (blue).

Following on from this, in order to gain deeper insight into gene level signal, we assessed the signal within genes that showed strongest support for just one of the three topologies. To do this, we carried out Approximately Unbiased (AU) tests (Shimodaira 2002) to quantify gene trees with enough phylogenetic signal to significantly reject all but one of the tested topologies. In theory, enriching for putative true orthologs should increase support for a single phylogeny. We wished to determine empirically whether this is the case. If hidden paralogy is present, we would expect that the CC fail datasets would have equal support for more than one topology while the CC pass set would support a single topology (Siu-Ting et al. 2019). However, we observed little or no conflict between CC fail and CC pass datasets in the subset of genes with the ability to reject all but one topology (Figure 5). The 46 gene trees in the Rouse2016 dataset capable of rejecting all but one topology (42 in CC fail, 4 in CC pass) show predominate support for T1 (i.e. Nephrozoa hypothesis), with 37/42 (88%) of the CC fail dataset supporting T1, and all 4 supporting T1 in the CC pass dataset. Of the 12 gene trees in the Cannon2016 dataset which were found to support a single topology (11 in CC fail, 1 in CC pass), we find 100% support for T1 in both the CC fail and CC pass datasets. For the 41 gene trees in the Philippe2019 dataset (37 in CC fail, 4 in CC pass) we find the same pattern with 100% support, but this time showing support for T3 (i.e. Xenambulacraria and paraphyletic Deuterostomia).

**Figure 5:**
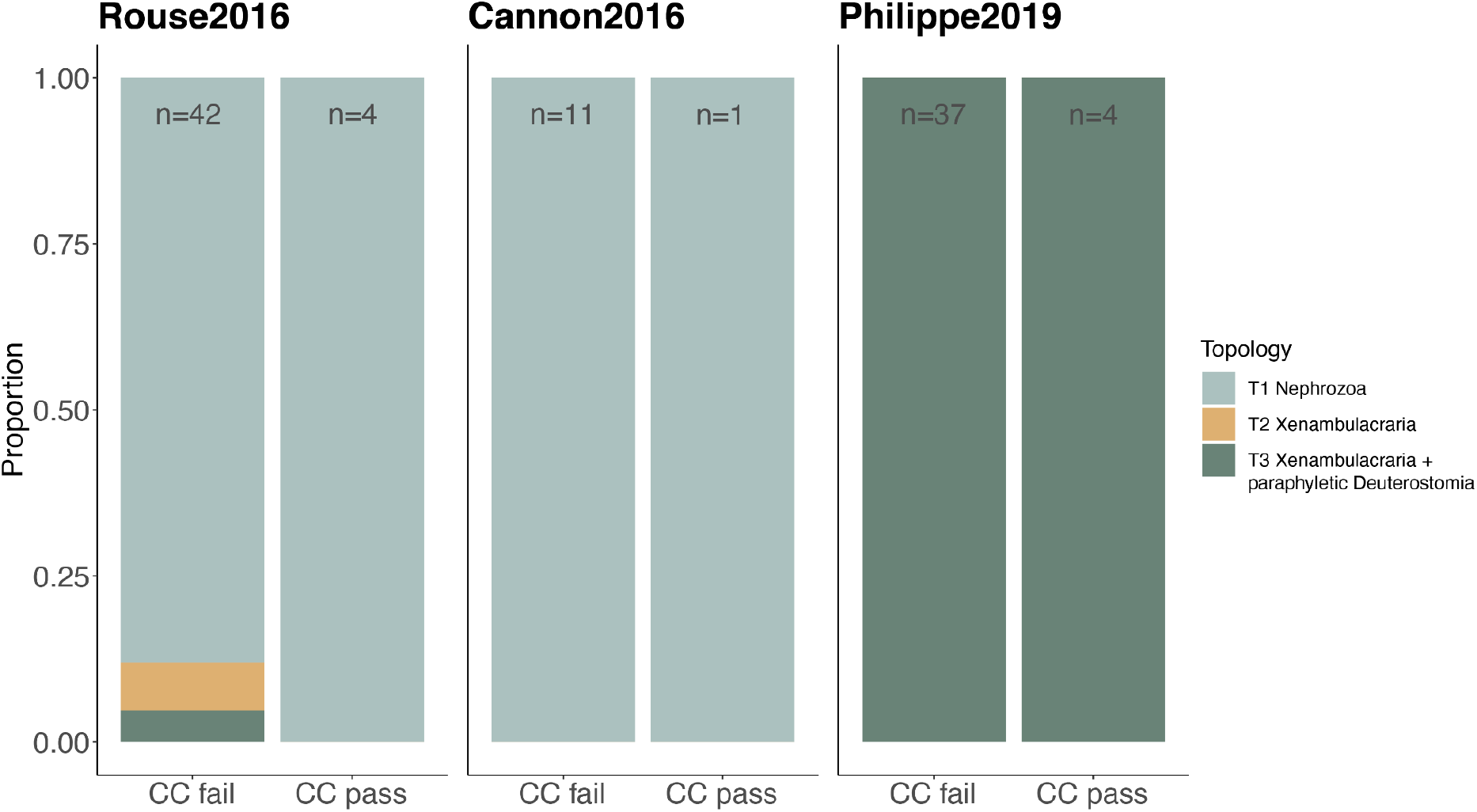
Distribution of signal in genes which reject all but one of the topologies: Support for alternative topologies in gene trees based on AU test. The x-axis contains columns for genes from CC fail and CC pass gene sets for each of the three datasets, Rouse2016, Cannon2016 and Philippe2019. The y-axis shows the proportion of genes supporting one of the three alternative topologies for the placement of Xenacoelomorpha. The number of genes in each subset (n) are shown for each of the three datasets.

### Increased support for Xenambulacraria hypothesis using genes enriched for orthology

Next, we carried out phylogenomic analyses on the filtered datasets using an ortholog enriched dataset, *i.e*. just the genes from the CC pass set, to determine the placement of Xenacoelomorpha. Our three CC pass subsets of Rouse2016, Cannon2016 and Philippe2019, consisted of 70, 16, and 65 genes respectively. These represented a total of 27,183, 4,080, and 27,448 aligned amino acid sites, significantly reduced compared to the original studies (Table 1). Phylogenomic reconstruction was carried out on each supermatrix using PhyloBayes-MPI (Lartillot et al. 2013) applying the CAT-GTR model, along a gamma distribution consisting of four rate categories (CAT-GTR+G4). Two independent chains were run for at least 10,000 iterations and convergence in all cases was assessed using the bpcomp function in PhyloBayes. All three analyses reached convergence between chains (with observed maxdiff < 0.3).

The CC pass subset of Rouse2016 recovered a topology consistent with the Xenambulacraria hypothesis (T2), with Xenacoelomorpha placed sister to Ambulacraria with high support (PP=0.94) (Figure 6, Supplementary figure S4). Deuterostomia was recovered as monophyletic, with a clade grouping Xenambulacraria + Chordata, albeit with lower support (PP=0.7). These findings conflict with the original study, where Xenacoelomorpha was sister to the remaining bilaterian lineages (Rouse et al. 2016), and instead supports grouping of Xenacoelomorpha within Deuterostomia, as found in in other previous studies (Philippe et al. 2011; Philippe et al. 2019). The CC pass subset of Cannon2016 supported the Nephrozoa hypothesis (T1), with Xenacoelomorpha sister to the remaining bilaterian lineages (PP=1) (Figure 6, Supplementary figure S5). Deuterostomia was found to be paraphyletic, with Chordata sister to Ambulacraria + Protostomia, however, with lower support (PP=0.81). This topology is in agreement with the placement of Xenacoelomorpha in the original study, but it does highlight the low support found for Deuterostomia monophyly (Philippe et al. 2019; Kapli et al. 2021). And finally, the CC pass subset of Philippe2019 supports the Xenambulacraria hypothesis placing Xenacoelomorpha sister to Ambulacraria (Xenambulacraria hypothesis) (PP=1.0) along with a paraphyletic Deuterostomia (PP=1.0) (T3) (Figure 6, Supplementary figure S6). In the original study, Philippe et al. (2019) proposed that Deuterostomia is non-monophyletic, although their support values were low. The analysis of the ortholog-enriched Philippe2019 data has also recovered non-monophyletic Deuterostomia, this time with full support (PP=1.0).

**Figure 6:**
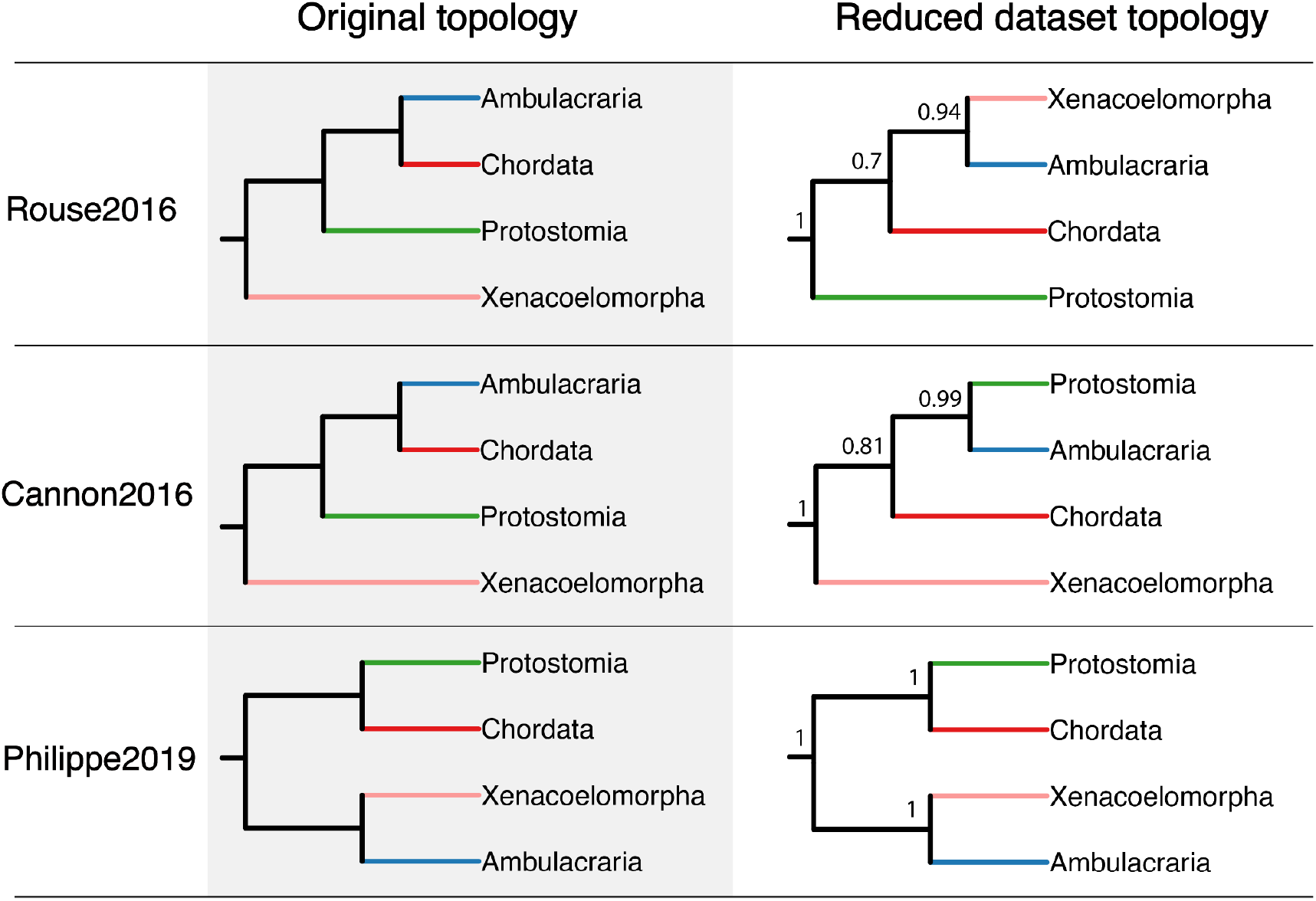
Phylogenetic trees generated from the CC Pass subsets of Rouse2016, Cannon2016, and Philippe2019. Original topologies are shown on the left in simplified form showing just the major bilaterian groups. The species topology from the ortholog enriched (i.e. CC Pass) subsets of each dataset are shown on the right. Posterior probabilities (PP) are shown for the nodes of major lineages.

### Filtering for improved orthology identification improves model fit

We applied posterior predictive analysis (PPA) to test for the adequacy of fit of the model to each of the three dataset (Bollback 2002). We used PhyloBayes-MPI (Lartillot et al. 2013) to infer five statistics (see materials and methods) designed to test model fit, which include three tests to describe among-site amino acid preferences as well as two to describe among-lineage compositional heterogeneity (Supplementary figure S7). In each case Z scores were used to test deviations for each test statistic from the null hypothesis, where a score of < 2 indicates that the model fits the data, while a Z score of > 5 indicates strong rejection of the null hypothesis that the model is capable of adequately describing the data (Feuda et al. 2017; Lartillot 2020). Analysing the results of the main test to measure whether the model adequately describes site specific amino acid preferences, PPA-DIV (Lartillot et al. 2007), we retrieve the following Z scores: Z < 2 for the Rouse2016 dataset indicating appropriate model fit in this statistic, and Z = 3.39 for Cannon2016 and Z=2.35 for Philippe2019 respectively (Table 2, Supplementary figure S7).

**Table 2.**
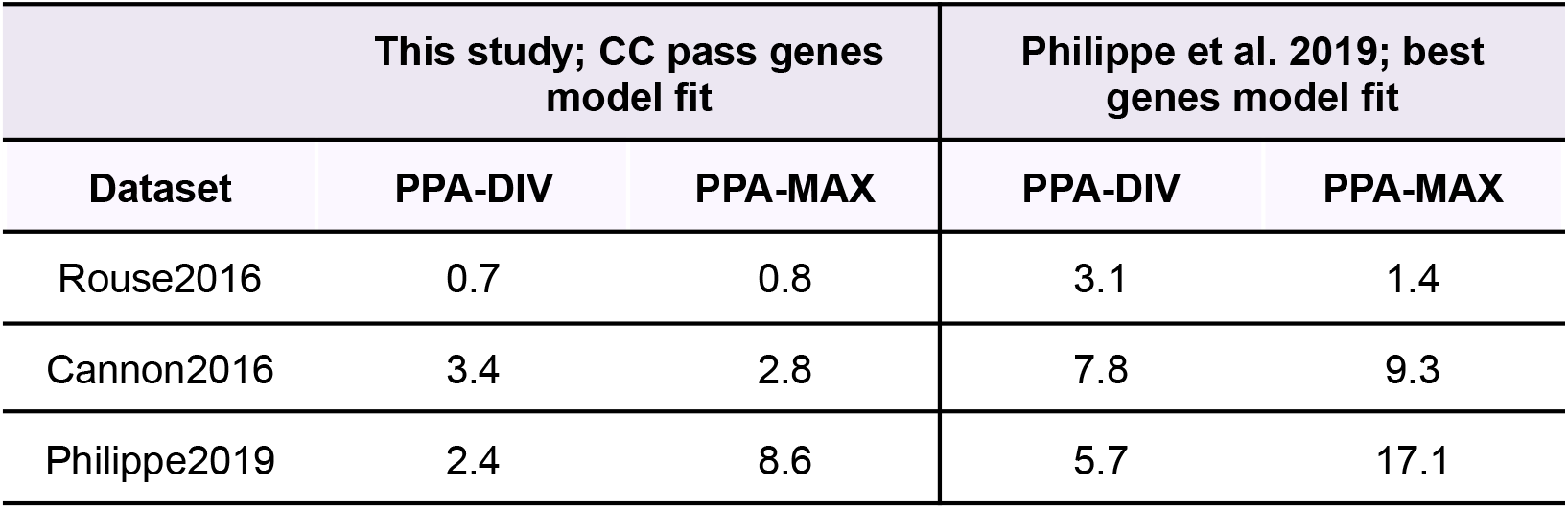
Comparing model fit between datasets. Comparison of the adequacy of fit of the CATGTR model to each of the three datasets. The left-most portion of the table refers to the Z scores for the CC pass gene sets constructed in this analysis for each of the three datasets. The two rightmost columns refer to analysis of Rouse2016 and Cannon2016 carried out by Philippe et al. (2019) where they compared model fit between the three datasets using a set of ‘best genes’ that contained an adequate amount of phylogenetic signal within their gene trees to be deemed useful for phylogenomic reconstruction. Each dataset shows the Z scores for PPA-DIV and PPA-MAX metrics of model fit.

The main test of among-lineage compositional heterogeneity, PPA-MAX, shows similar patterns, with a Z score of < 2 for Rouse2016 dataset, and Z scores > 2 for both Cannon2016 (2.77) and Philippe2019 (8.61) datasets (Table 2, Supplementary figure S4). The Z score of greater than 5 for the reduced Philippe2019 dataset suggests that the model is not adequately estimating the maximal compositional heterogeneity observed across taxa in the dataset. This may be due to the fact that in the original analysis by Philippe et al. (2019) the genes were not filtered for variation in rates of compositional heterogeneity, while in the original study by Cannon et al. (2016) a filter was applied to account for this (although there was no filter of the data for such patterns by Rouse et al. (2016), the low Z score may be due to the smaller number of species in this dataset). The only other study to test for model fit using PPA was Philippe et al. (2019). In their analysis they compared the model fit in the best and worst genes (based on a metric for phylogenetic signal within the gene tree) between the CAT and CATGTR models in each of the three datasets we have used here (Philippe et al. 2019). While they observed improved model fit, determined by the Z score from PPA-DIV, PPA-MAX and PPA-MEAN, in the best genes applying the CATGTR model, the Z scores observed in our reduced datasets show further improvement in Z scores suggesting increased model fit (Table 2, Supplementary figure S4). However, while we do see an increase in model fit, these metrics suggest that the CATGTR model does not adequately model the evolutionary processes in each of the three datasets.

## Discussion

Where Xenacoelomorpha place in the animal tree of life has profound implications for our understanding of animal evolution. The patterns in the data previously applied to this problem indicate low levels of signal (Figure 2), suggesting a small proportion of genes with strong signal which are contributing to the placement of Xenacoelomorpha (Shen et al. 2017; Brown and Thomson 2017; Di Franco et al. 2019; Walker et al. 2020). The issue of inadequate or misleading signal within phylogenomic studies is often overlooked in favour of appropriate model selection or the size of data matrices (Philippe, Brinkmann, Lavrov, et al. 2011). The work presented here and elsewhere (Shen et al. 2017; Siu-Ting et al. 2019; Francis & Canfield 2020) demonstrates that without assessment of the underlying molecular data, we risk misinterpreting the signal present (Brown & Thomson 2017; Shen et al. 2017; Siu-Ting et al. 2019; Francis & Canfield 2020). Our approach to this problem is to reduce the data matrices to only include genes enriched for orthology. While inferring the ‘true’ species tree is still difficult with a filtered dataset, accounting for and reducing genes with potentially misleading signal does provide a sound criteria for species tree inference (Salichos & Rokas 2011; Doyle et al. 2015; Edwards 2016; Shen et al. 2016, 2017; Brown & Thomson 2017; Kocot et al. 2017; Molloy & Warnow 2018; Dornburg et al. 2019; Siu-Ting et al. 2019; Smith et al. 2020; Koch 2021). We have shown that in most cases reducing the dataset down to genes that can recapitulate known splits in the animal tree increases overall quality of signal while allowing for improved modelling of the data. It could therefore be argued that reducing datasets in this way generates datasets with as much, if not more, power of resolution than their larger alternatives (Koch 2021). Importantly, these reduced datasets have improved model fit and tractable time scales for reaching convergence in Bayesian phylogenomic analyses. However, we show that while the fit of the model to the data is improved, there is still the issue of model inadequacy even when applying highly parameterised models like CAT-GTR on these reduced data matrices. Future studies would benefit from development of models which can better accommodate the heterogeneity within the data, in particular compositional heterogeneity, in a parallelised way (Foster 2004).

The question of whether it is hidden paralogy alone driving these misleading effects in species tree inference also requires further investigation. The consistent support for a single topology from CC pass and CC fail gene sets provides additional support for other sources of misleading signal at play (Siu-Ting et al. 2019; Shen et al. 2021; Smith & Hahn 2021). Using simulations it has been demonstrated that Clan_Check can account for ILS (Siu-Ting et al. 2019), therefore, while inadvertent paralog inclusion is an issue, more work is needed to tease apart other potential hidden factors causing gene tree discordance. After filtering for biases within the data, the influence of dataset size is another aspect which requires further attention. There were only 16 genes which passed the filtering step for Cannon2016, which perhaps unsurprisingly resulted in polytomies in the certain parts of the topology (i.e. between Placozoa, Cnidaria and Bilatera). Caution is recommended when reducing data matrices to such small sizes as there may be inadvertent effects due to low levels of total phylogenetic signal. Shifts in the placement of Xenacoelomorpha following reanalysis of these previously published datasets (Rouse et al. 2016; Cannon et al. 2016) were also reported by Philippe et al. (2019). However, our approach using a subset of genes filtered for paralogs, provides a species topology with greater overall support in all cases as well as improvement in model fit to the data.

We find increased, but not complete, support in our reduced datasets for the Xenambulacraria clade (Figure 6). We also find increased but not consistent support throughout our analyses for the paraphyly of Deuterostomia. This lack of signal for the existence of Deuterostomia is also reflected in the very small number of genes which recapitulate this node (Figure 2A). It is clear that studies focussed on the placement of Xenacoelomorpha, and other challenging nodes, would benefit from tractable and consistent methods, as well as from greater exploration of the signal within the data matrices. Currently, there is a clear deficit in the signal to noise ratio in our phylogenomic studies assembled to resolve this position. Here, we attempted to increase the strength of true phylogenetic signal by using only genes which we know can recapitulate uncontroversial splits and have succeeded in generating datasets enriched for orthology that are modeled more adequately. Moving forward, the generation of more complete genomes for groups such as Xenacoelomorpha, is of key strategic importance, as is consilience across alternative data types such as rare genomic changes (Rokas & Holland 2000).

## Acknowledgements

MJO’C would like to thank both the University of Leeds for her 250 Great Minds Fellowship career award and the University of Nottingham for funding this work through POM and CGPM. This work was undertaken on ARC3, part of the High Performance Computing facilities at the University of Leeds, UK. C.J.C. wishes to acknowledge funding from the BBSRC (BB/R019185/1) and DAFM Ireland/DAERA Northern Ireland (R3192GFS) and both C.J.C and K.S.-T. from the EC, Horizon 2020 (818368, MASTER). This work was supported by a fellowship from the Irish Research Council–Marie Sklodowska-Curie cofund program (ELEVATEPD/2014/69 to K.S.-T.);

## Author contributions

MJO’C conceived the project and directed the work. POM and CGPM carried out all analyses with input from MJO’C. KST and CJC contributed to the experimental design. MJO’C and POM wrote the manuscript and all authors contributed to editing.

## Methods

### Measuring rate of violation of known monophyletic clans with Clan_Check

Our aim in this study is to test whether hidden paralogy in phylogenomic datasets, as a result of ancient gene duplication followed by loss, is driving alternative topologies in the placement of Xenacoelomorpha. To do this we employed Clan_Check (https://github.com/ChrisCreevey/clan_check) which is a tool designed to test if assumed single copy orthologous gene families violate incontestable monophyletic clans, as a result of hidden paralogy (Siu-Ting et al. 2019). Phylogenomic datasets for all three studies were downloaded from online data depositories: datadryad.org/stash/dataset/doi:10.5061/dryad.79dq1 (Rouse et al. 2016), datadryad.org/stash/dataset/doi:10.5061/dryad.493b7, (Cannon et al. 2016) and github.com/MaxTelford/Xenacoelomorpha2019 (Philippe et al. 2019). These datasets are referred to as Rouse2016, Cannon2016 and Philippe2019 throughout the main text. Each dataset consisted of a concatenated matrix of each of their gene families in amino acid format (see Table 1 for details on the number of gene families and species in each dataset). This concatenated matrix was split into constituent gene family fasta files and each one was aligned using three alignment software methods; Mafft (Katoh et al. 2005), Muscle (Edgar 2004) and Prank (Löytynoja & Goldman 2005). In order to remove any potential bias on downstream analyses from the alignment step, we selected the best alignment using MetAl (Blackburne & Whelan 2012). MetAl calculates metric distances between alignments of the same sequence, where a score of < 0.15 between a pair of alignments were considered to be in agreement while a score of > 0.15 were considered discordant. If a pair of alignments were found to be discordant, alignment quality was assessed using norMD and the alignment with the highest similarity score was retained for subsequent tree inference analyses (Thompson et al. 2001; Muller et al. 2010). If there was no alignment with a greater norMD score than the other two, the Mafft gene alignment was selected. Next, we used IQ-TREE (Nguyen et al. 2015), applying ModelFinder (Kalyaanamoorthy et al. 2017) to find the model of best fit and carrying out 1000 ultrafast bootstrap replicates to construct gene trees for all gene family alignments in each dataset. For each dataset, we annotated a number of clans which were to be tested using Clan_Check. These were assigned based on the phylogenetic spread of species within each of the dataset. The clans/splits assigned for testing included Porifera, Ctenophora, Cnidaria, Bilateria, Protostomia, Deuterostomia, Xenacoelomorpha, Ambulacraria, Lophotrochozoa, Ecdysozoa, and Chordata (Supplementary Figure S2). Monophyly of Ctenophora was only tested on Cannon2016, as there was just one species sampled for this clade in Rouse2016 and no species for Philippe2019. For a gene family to pass the Clan_Check filter, we required the corresponding gene tree to recapitulate a given number of clans. For Rouse2016 and Philippe2019 this cut off was 4 and 5 clans, respectively. For the Cannon2016 dataset, the cut off was set to 3 clans. This was due to the fact that none of the gene trees in this dataset could recapitulate 5 clans, and only 4 gene trees could recapitulate 4 clans (Supplementary Table S1). Using these dataset specific cutoffs retained a similar proportion of the overall number orthogroups for each of the reduced datasets.

### Metrics to test for biases and signal between CC pass and CC fail datasets

For each of the three datasets we compared the sets of genes that passed and failed the Clan_Check filter step, first to ensure that the sampled set of genes for downstream analyses were not biased towards branch length, compositional heterogeneity or function. Here we followed the protocol used in Siu-Ting et al (2019). To check for bias in branch length between the paralogous or orthologous gene sets, we used the inferred gene trees to compare distributions of branch lengths between the two sets. Average branch lengths were calculated by summing all branch lengths in the gene tree and dividing by the total number of branches. A Wilcox signed-rank test was then run in R (R Core Team 2013) to check whether there was a significant difference in average branch lengths between the two datasets. We used the same statistical test to check whether there were differences between the compositional heterogeneity of the taxa sampled in each dataset. We generated gene trees for each of the alignments and obtained the proportion of taxa that passed or failed the compositional heterogeneity test in IQTree (Minh et al. 2020) after running each of the aligned gene families to infer a gene tree. Finally, to test whether there was a bias in the functions of genes which passed or failed the Clan_Check filter, we ran gene ontology analysis on the CC pass and CC fail gene sets. Gene annotation for each orthogroup was carried out using the representative human sequence, where possible, using InterProScan (Jones et al. 2014). Gene ontologies were then extracted for each orthogroup and were grouped into one of the three major GO categories; “Molecular Function”, “Biological Process” or “Cellular Component”. This hierarchical annotation of GO terms was carried out using the GOATools python library (Klopfenstein et al. 2018). Finally, we ran a Pearson’s chi-squared test between the putative orthologous and paralogous gene sets to test whether either of the gene sets had statistically significant different distribution of functional categories.

We also ran a number of gene and tree based metrics to compare the level of phylogenetic signal between the two gene sets. This was to ensure that the gene families enriched for orthologous genes actually displayed greater levels of phylogenetic signal than those that were filtered out. In total seven sets of tests were applied to the pass and fail gene sets for each of the three datasets. These included alignment length, bipartition support, long branch score, number of parsimony informative sites, level or saturation, treeness divided by relative compositional variability (RCV), and the number of variable sites. Each of these tests were measured using the PhyKIT software (Steenwyk et al. 2021). Briefly, alignment length has been shown to correlate with accurate tree inference (Shen et al. 2016), gene trees which display higher bipartition support have been found to display greater certainty among bipartitions (Salichos and Rokas 2013; Shen et al. 2016), a larger number of parsimony informative sites is associated with stronger phylogenetic signal (Shen et al. 2016; Steenwyk et al. 2020), the average long branch score in a tree may give insights into the level of heterogeneity within the gene (Struck 2014), saturation is driven by sites with multiple substitutions thus underestimating the genetic distance among taxa, with a score of 1 showing no saturation and a score of 0 showing complete saturation, genes with higher treeness divided by RCV has been found to be associated with lower compositional and other biases and higher phylogenetic signal (Philippe, Brinkmann, Lavrov, et al. 2011), and finally genes with a higher number of variable sites often display higher phylogenetic signal (Shen et al. 2016).

### Testing gene family support for alternative topologies between pass and fail datasets

To examine the level of support for each of the three alternative topologies for the placement of Xenacoelomorpha (T1, T2, and T3), we carried out two sets of analyses. First, we ran a gene-wise likelihood test, as in Shen et al. (2017), to measure the distribution of signal for each of the three topologies between the genes that failed and pass the Clan_Check filter. We constructed constrained species trees based on each of the three alternative topologies. Then, site-wise log likelihood estimations were inferred by comparing the data matrix of genes to each alternative topology using RAxML and Phylogenetic_signal_parser perl script from Shen et al. (2017) (https://figshare.com/articles/dataset/Contentious_relationships_in_phylogenomic_studies_can_be_driven_by_a_handful_of_genes/3792189/4).

Secondly, we ran AU tests on gene trees for the pass and fail datasets to calculate the number of genes capable of statistically significantly rejecting all but one of three alternative topologies. First, we constructed idealised gene trees consistent with each of the three alternative topologies for each dataset using Clann (Creevey & McInerney 2005). IQTree was used to construct a ML tree for each of the gene family alignments, and then calculate the log-likelihood of each of the three alternative species tree topologies based on the estimated parameters inferred for the ML gene tree. For each alternative topology the AU test returns a p-value, where a tree is rejected with a p-value < 0.05. Thus, if a gene tree can confidently reject all but one of the alternative topologies, the topology supported was recorded. Again, this was carried out to compare between the gene sets that passed and failed the Clan_Check filter.

### Phylogenomic analyses using reduced datasets

After filtering each of the genesets from the three studies using Clan_Check, concatenated matrices of aligned amino acid sequences in Phylip format were constructed using SCaFOs and TREE-PUZZLE with default options (Schmidt et al. 2002; Roure et al. 2007). For each resulting supermatrix, phylogenetic reconstruction was carried out using PhyloBayes-MPI (Lartillot et al. 2013). After constant sites were removed (-dc option) the CAT-GTR model was applied, along a gamma distribution consisting of four rate categories. Two independent chains were run until convergence between the runs was reached. Convergence between chains was assessed using the bpcomp function in PhyloBayes with a burn-in of 5,000 iterations and sampling every 10 iterations. A maxdiff score of below 0.3 indicated convergence. Trees were visualised using the ggtree package in R (Supplementary Figures S4, S5 and S6)(Yu et al. 2017).

### Posterior predictive analysis to assess model fit

Posterior predictive analysis (PPA) was applied in PhyloBayes-MPI to assess how well the CAT-GTR model fit each of the reduced datasets (Bollback, 2002; Lartillot and Philippe, 2004; Lartillot et al., 2013; Feuda et al., 2017). The allppred flag in the readpb_mpi module was used to perform PPAs on both chains used for phylogenomic reconstruction. We tested five statistics to measure model fit, including three for assessing site-specific heterogeneity (PPA-DIV, PPA-CONV and PPA-VAR) and two for lineage-specific heterogeneity (PPA-MAX and PPA-MEAN). For each statistic, a Z-score was computed with |Z| representing standard deviations of the simulated data from the observed mean for each statistic (Feuda et al., 2017). The resulting Z scores were then compared to those calculated for the CAT-GTR model in Philippe et al. (2019).

### Data and code availability

The raw fasta files, alignments, gene trees and species trees presented in this study, along with all code required to reproduce each analysis, are available from GitHub (https://github.com/PeterMulhair/Xenaceol_Paralogy).

## Supplementary files

### Supplementary Figures

**Supplementary Figure S1:**
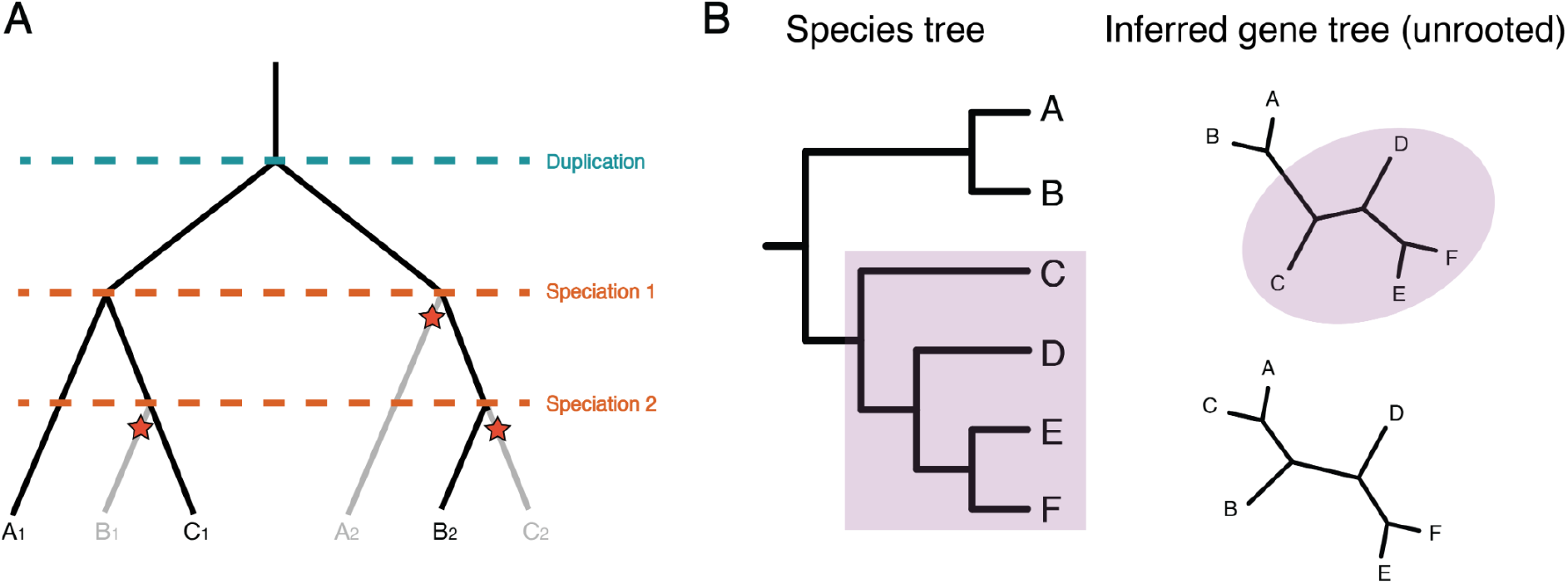
Summary of concepts behind Clan_Check analysis. (A) Gene tree which represents the evolution of a single example gene in three species (A, B, and C). Ancient duplication of the gene (Duplication) is followed by two rounds of speciation events (Speciation 1 and Speciation 2), resulting in two gene copies in each species eg. species A has gene A1 and A2. Subsequent loss of gene copies are labelled with red stars. This results in a mix of ortholog and paralog genes within the extant species. The species relationships should be given as species B sister to species C, with species A as the outgroup to this clade. However, in this example, the hidden paralog B2 which is a result of the late loss of ortholog and paralog genes causes species A to group with C, to the exclusion of B. (B) Summary of Clan_Check test where known monophyletic groups are defined from the species tree (eg. clade consisting of C, D, E and F). Gene trees are then inferred and the presence of the clan containing these species are tested. Top gene tree is able to recapitulate the clan consisting of the four taxa, whereas the bottom gene tree violates our assumptions of the species tree with these taxa not forming a monophyletic clan.

**Supplementary Figure S2:**
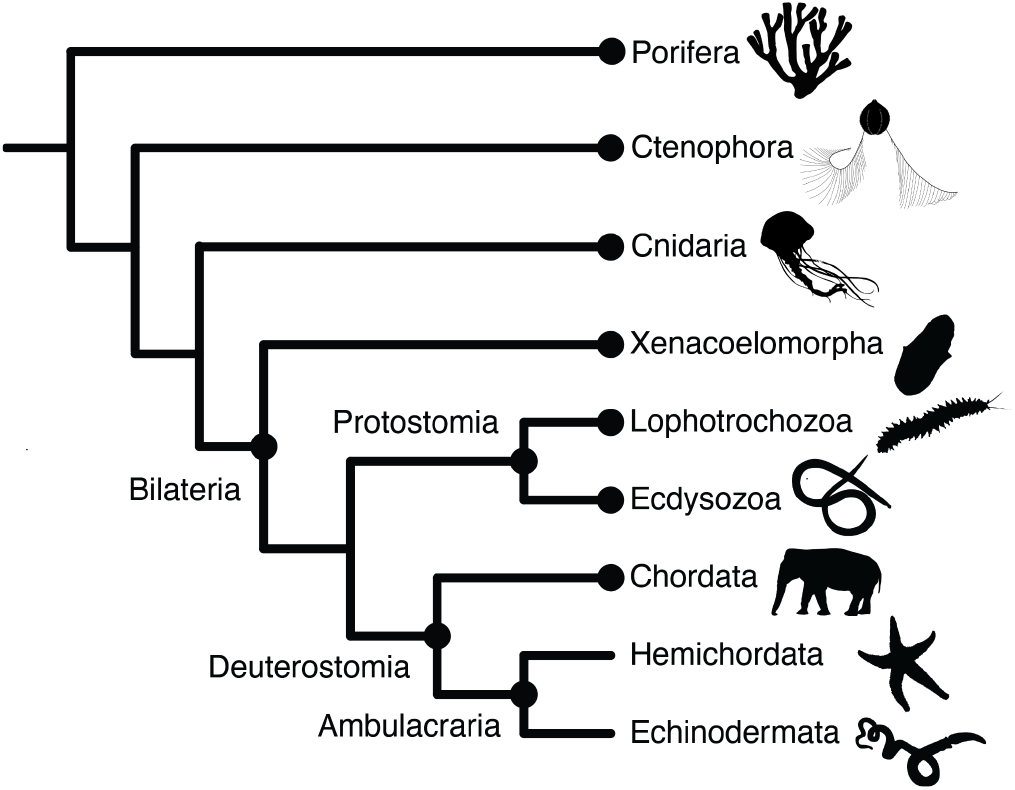
Clans on the animal tree whose monophyly were tested in each of the gene trees using Clan_Check. The clans tested are labelled with a dark circle and the name of the clan.

**Supplementary Figure S3:**
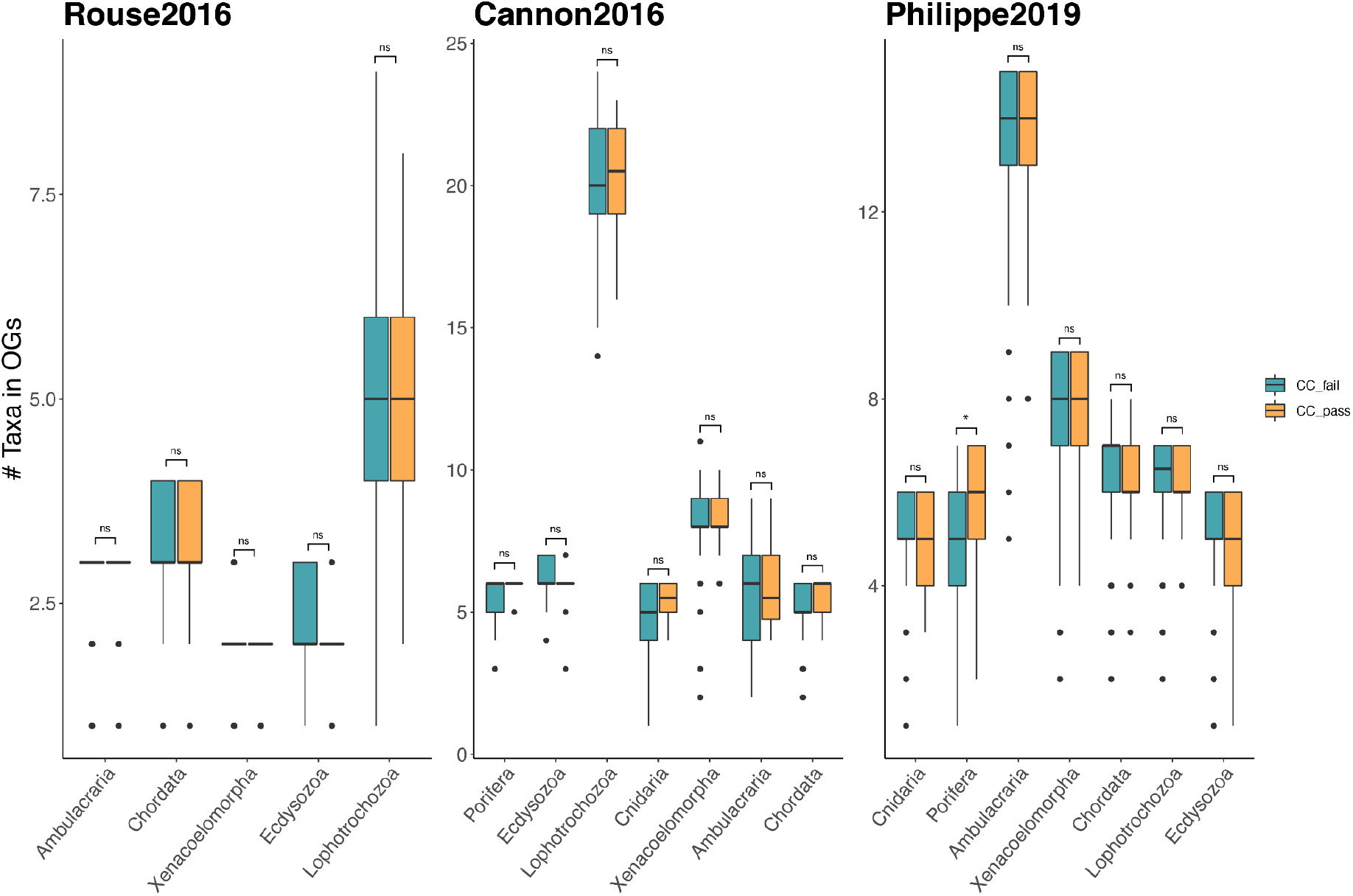
Check for changes in taxon sampling between CC fail and CC pass gene sets. For each dataset we plotted the number of species (y-axis) in each phyla (x-axis) in each OG in the CC fail and CC pass gene sets. (NS = *P*-value > 0.05, \**P*-value < 0.05, \*\**P*-value < 0.01, \*\*\**P*-value < 0.001).

**Supplementary Figure S4:**
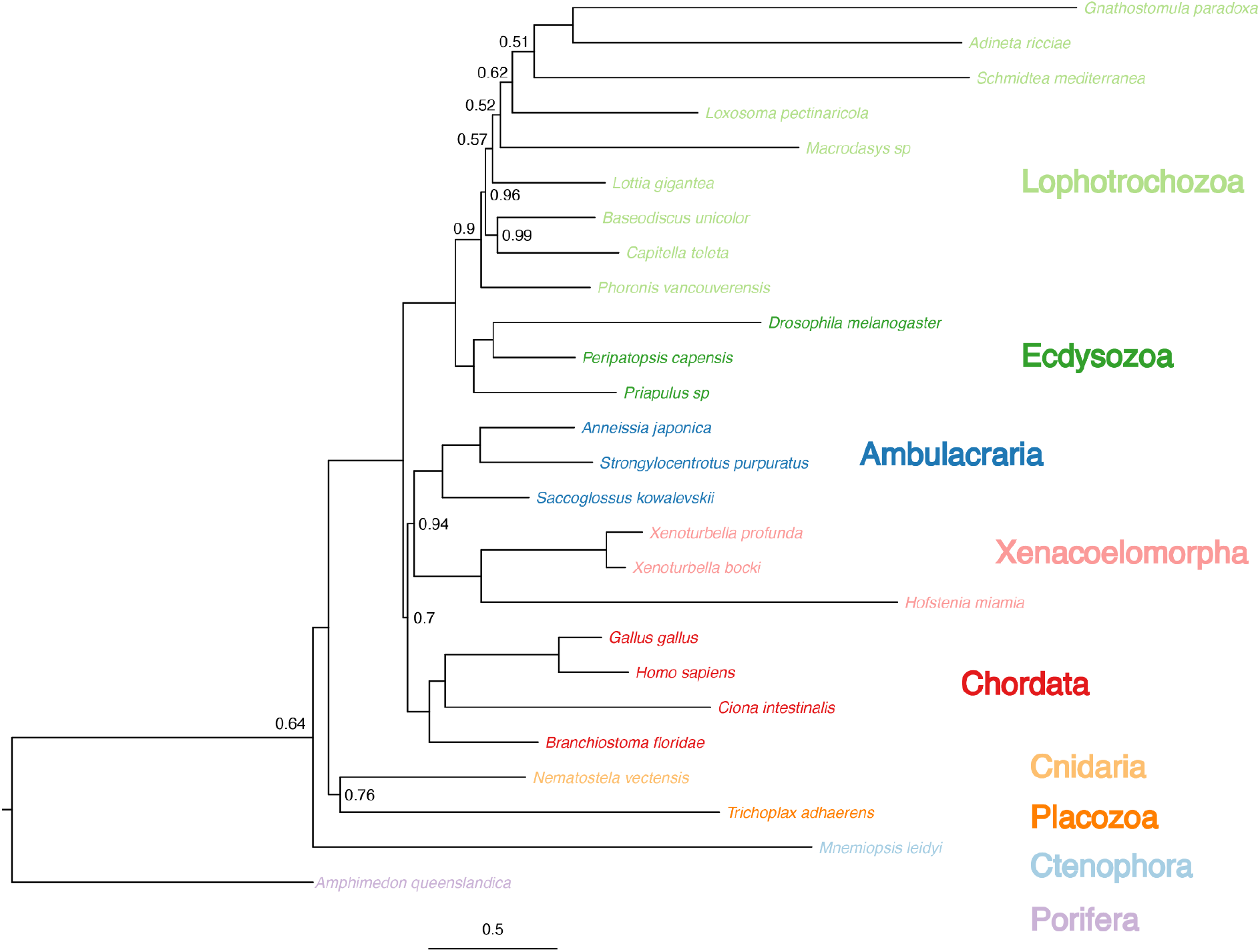
Species trees inferred from reduced CC pass data matrix for the Rouse2016 dataset. Bayesian species tree inferred using the reduced dataset of 70 genes, and the CAT-GTR+G4 model. Major clades are coloured per lineage, and node support, where PP is below 1, is annotated on the tree. Xenacoelomorpha are grouped with Ambulacraria, with this clade sister to Chordata.

**Supplementary Figure S5:**
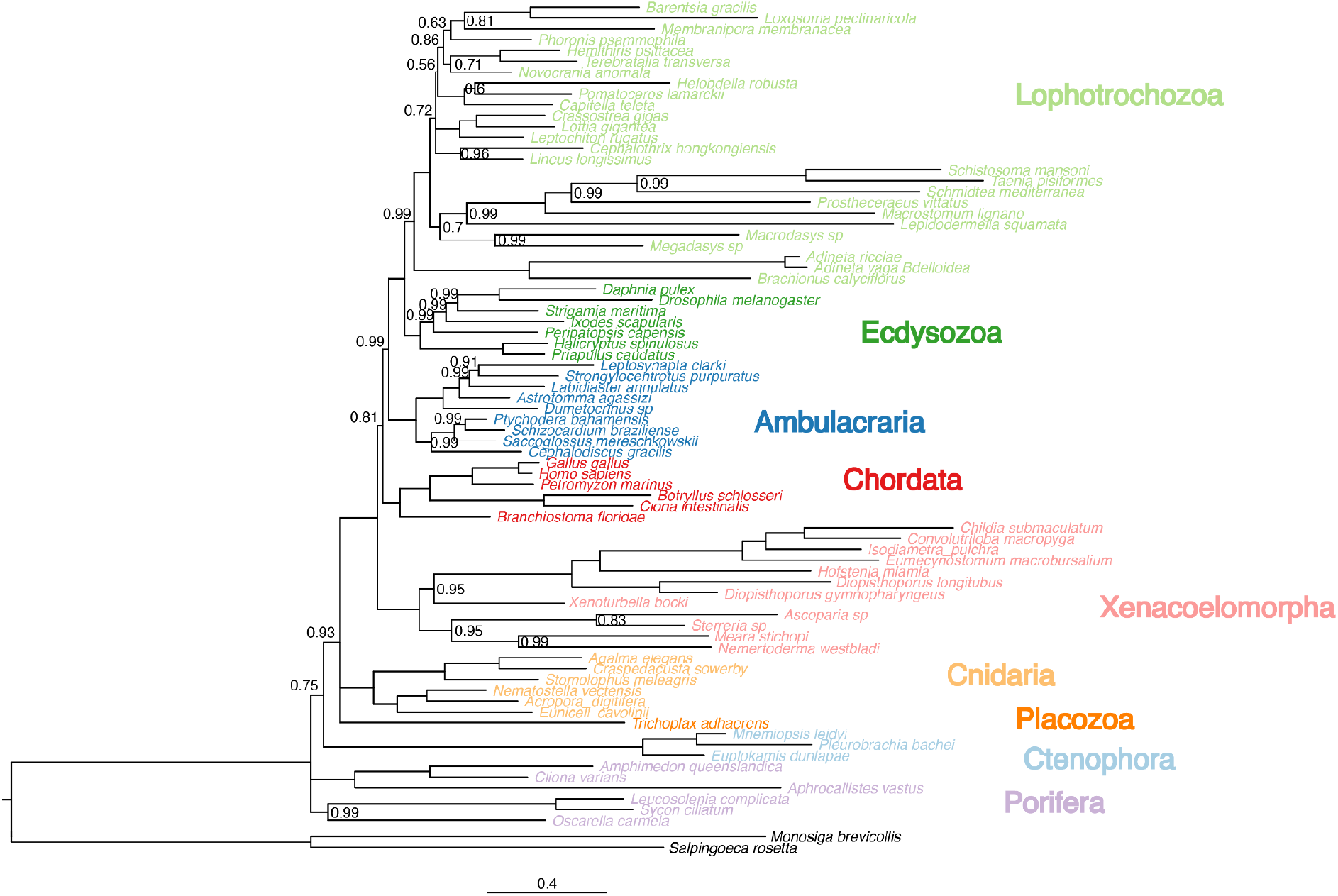
Species trees inferred from reduced CC pass data matrix for the Cannon2016 dataset. Bayesian species tree inferred using the reduced dataset of 16 genes, and the CAT-GTR+G4 model. Major clades are coloured per lineage, and node support, where PP is below 1, is annotated on the tree. Xenacoelomorpha are placed sister to the remaining bilaterian groups.

**Supplementary Figure S6:**
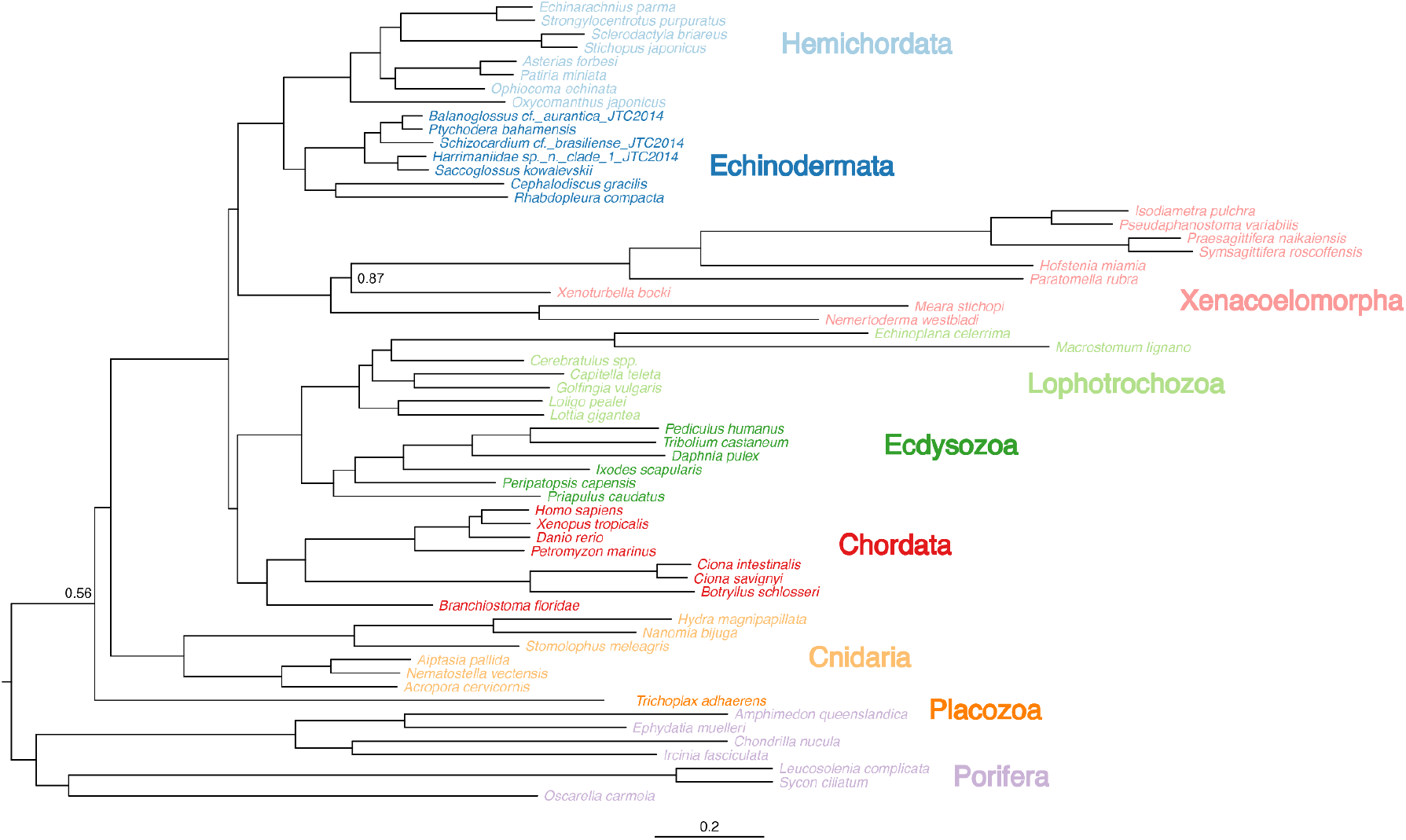
Species trees inferred from reduced CC pass data matrix for the Philippe2019 dataset. Bayesian species tree inferred using the reduced dataset of 65 genes, and the CAT-GTR+G4 model. Major clades are coloured per lineage, and node support, where PP is below 1, is annotated on the tree. Xenacoelomorpha are placed sister to Ambulacraria. Chordata group with the two Protostomia phyla implying paraphyletic Deuterostomia.

**Supplementary Figure S7:**
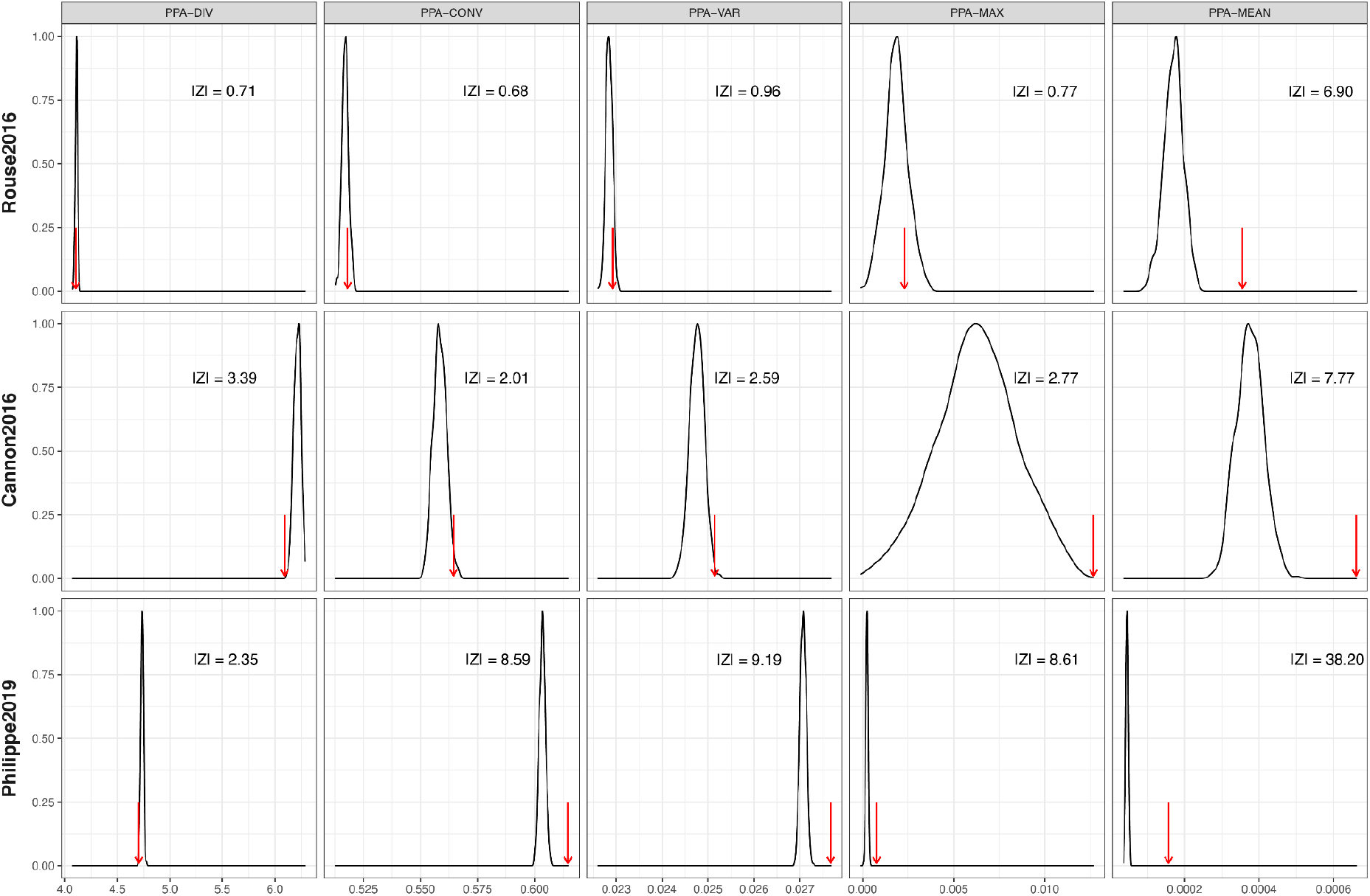
Model fit assessment for each of the three filtered datasets (Rouse2016, Cannon2016 and Philippe2019). Each column shows one of five posterior predictive analysis (PPA) statistics; from left to right, PPA-DIV, PPA-CONV, PPA-VAR, PPA-MAX, and PPA-MEAN. Each red row represents model fit metrics for each of the three filtered datasets. The Z score (|Z|) represent deviations from the null hypothesis for each test statistic.

### Supplementary Tables

**Supplementary Table S1:**
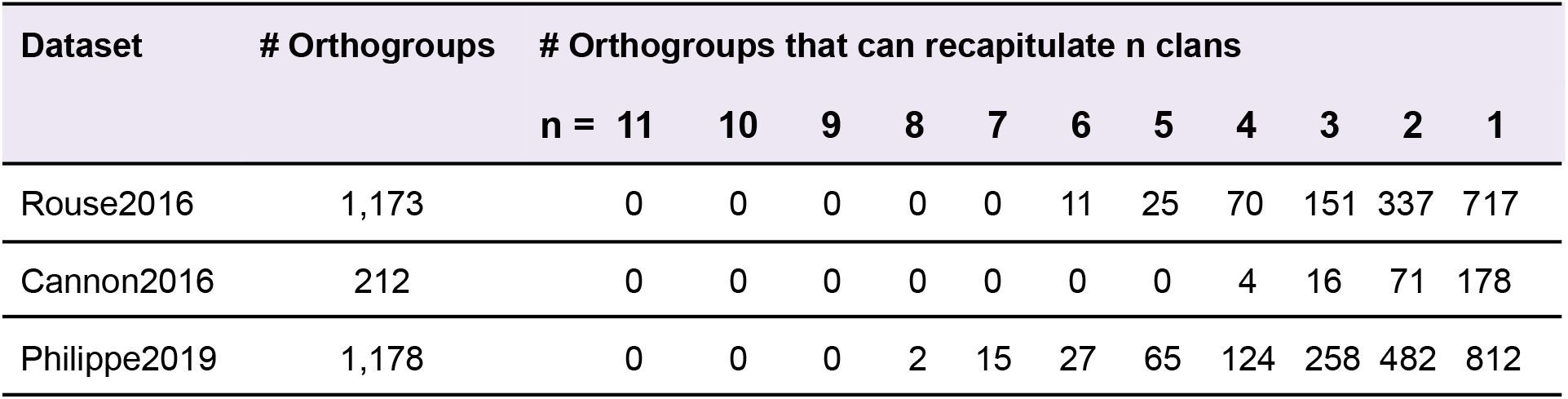
Number of clans recapitulated by gene trees for each of the three previously published datasets. For each dataset, the number of orthogroups are given, for each a gene tree was constructed. The presence of 11 incontestable clans were then tested in each gene tree. The far right column represents the number of genes which could recapitulate a number of these clans, in descending order.

**Supplementary Table S2:**
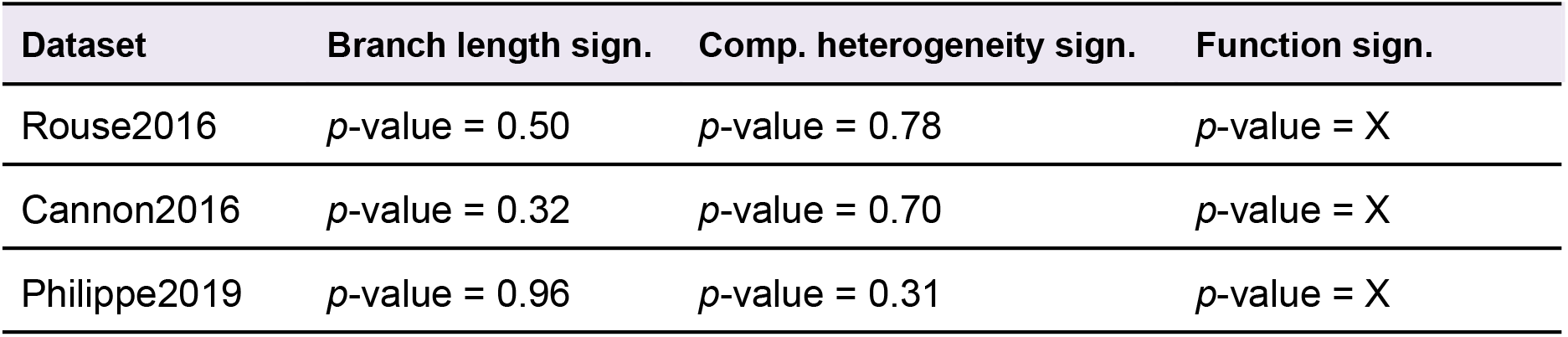
Results of tests for statistical significance between CC pass and CC fail gene sets for branch length, compositional heterogeneity and function. For each dataset, a Wilcoxon rank-sum test was carried out between the CC pass and CC fail gene sets for each of the three metrics annotated. *P*-value of < 0.05 was considered significant.

## References

Altenhoff AM et al. 2019. OMA standalone: orthology inference among public and custom genomes and transcriptomes. Genome Res. 29:1152–1163.

Blackburne BP, Whelan S. 2012. Measuring the distance between multiple sequence alignments. Bioinformatics. 28:495–502.

Bollback JP. 2002. Bayesian model adequacy and choice in phylogenetics. Mol. Biol. Evol. 19:1171–1180.

Bourlat SJ et al. 2006. Deuterostome phylogeny reveals monophyletic chordates and the new phylum Xenoturbellida. Nature. 444:85–88.

Bourlat SJ, Nielsen C, Lockyer AE, Littlewood DTJ, Telford MJ. 2003. Xenoturbella is a deuterostome that eats molluscs. Nature. 424:925–928.

Brown JM, Thomson RC. 2017. Bayes Factors Unmask Highly Variable Information Content, Bias, and Extreme Influence in Phylogenomic Analyses. Syst. Biol. 66:517–530.

Cannon JT et al. 2016. Xenacoelomorpha is the sister group to Nephrozoa. Nature. 530:89–93.

Creevey CJ, McInerney JO. 2005. Clann: investigating phylogenetic information through supertree analyses. Bioinformatics. 21:390–392.

Di Franco A, Poujol R, Baurain D, Philippe H. 2019. Evaluating the usefulness of alignment filtering methods to reduce the impact of errors on evolutionary inferences. BMC Evol. Biol. 19:21.

Dornburg A, Su Z, Townsend JP. 2019. Optimal Rates for Phylogenetic Inference and Experimental Design in the Era of Genome-Scale Data Sets. Syst. Biol. 68:145–156.

Doyle VP, Young RE, Naylor GJP, Brown JM. 2015. Can We Identify Genes with Increased Phylogenetic Reliability? Syst. Biol. 64:824–837.

Dunn CW, Howison M, Zapata F. 2013. Agalma: an automated phylogenomics workflow. BMC Bioinformatics. 14:330.

Ebersberger I, Strauss S, von Haeseler A. 2009. HaMStR: profile hidden markov model based search for orthologs in ESTs. BMC Evol. Biol. 9:157.

Edgar RC. 2004. MUSCLE: a multiple sequence alignment method with reduced time and space complexity. BMC Bioinformatics. 5:113.

Edwards SV. 2016. Phylogenomic subsampling: a brief review. Zool. Scr. 45:63–74.

Feuda R et al. 2017. Improved Modeling of Compositional Heterogeneity Supports Sponges as Sister to All Other Animals. Curr. Biol. 27:3864–3870.e4.

Foster PG. 2004. Modeling compositional heterogeneity. Syst. Biol. 53:485–495.

Francis WR, Canfield DE. 2020. Very few sites can reshape the inferred phylogenetic tree. PeerJ. 8:e8865.

Haszprunar G. 2016. Review of data for a morphological look on Xenacoelomorpha (Bilateria incertae sedis). Org. Divers. Evol. 16:363–389.

Hejnol A, Pang K. 2016. Xenacoelomorpha’s significance for understanding bilaterian evolution. Curr. Opin. Genet. Dev. 39:48–54.

Jones P et al. 2014. InterProScan 5: genome-scale protein function classification. Bioinformatics. 30:1236–1240.

Kalyaanamoorthy S, Minh BQ, Wong TKF, von Haeseler A, Jermiin LS. 2017. ModelFinder: fast model selection for accurate phylogenetic estimates. Nat. Methods. 14:587–589.

Kapli P et al. 2021. Lack of support for Deuterostomia prompts reinterpretation of the first Bilateria. Science Advances. 7:eabe2741.

Kapli P, Telford MJ. 2020. Topology-dependent asymmetry in systematic errors affects phylogenetic placement of Ctenophora and Xenacoelomorpha. Science Advances. 6:eabc5162.

Katoh K, Kuma K-I, Toh H, Miyata T. 2005. MAFFT version 5: improvement in accuracy of multiple sequence alignment. Nucleic Acids Res. 33:511–518.

Klopfenstein DV et al. 2018. GOATOOLS: A Python library for Gene Ontology analyses. Sci. Rep. 8:10872.

Koch NM. 2021. Phylogenomic subsampling and the search for phylogenetically reliable loci. Mol. Biol. Evol. doi: 10.1093/molbev/msab151.

Kocot KM et al. 2017. Phylogenomics of Lophotrochozoa with Consideration of Systematic Error. Syst. Biol. 66:256–282.

Lartillot N. 2020. The bayesian approach to molecular phylogeny. https://hal.archives-ouvertes.fr/hal-02535330/file/chapter_1.4_lartillot_chap.pdf.

Lartillot N, Brinkmann H, Philippe H. 2007. Suppression of long-branch attraction artefacts in the animal phylogeny using a site-heterogeneous model. BMC Evol. Biol. 7 Suppl 1:S4.

Lartillot N, Rodrigue N, Stubbs D, Richer J. 2013. PhyloBayes MPI: phylogenetic reconstruction with infinite mixtures of profiles in a parallel environment. Syst. Biol. 62:611–615.

Löytynoja A, Goldman N. 2005. An algorithm for progressive multiple alignment of sequences with insertions. Proc. Natl. Acad. Sci. U. S. A. 102:10557–10562.

Minh BQ et al. 2020. IQ-TREE 2: New Models and Efficient Methods for Phylogenetic Inference in the Genomic Era. Mol. Biol. Evol. 37:1530–1534.

Molloy EK, Warnow T. 2018. To Include or Not to Include: The Impact of Gene Filtering on Species Tree Estimation Methods. Syst. Biol. 67:285–303.

Morgan CC et al. 2013. Heterogeneous models place the root of the placental mammal phylogeny. Mol. Biol. Evol. 30:2145–2156.

Muller J, Creevey CJ, Thompson JD, Arendt D, Bork P. 2010. AQUA: automated quality improvement for multiple sequence alignments. Bioinformatics. 26:263–265.

Paps J, Baguñá J, Riutort M. 2009. Bilaterian phylogeny: a broad sampling of 13 nuclear genes provides a new Lophotrochozoa phylogeny and supports a paraphyletic basal acoelomorpha. Mol. Biol. Evol. 26:2397–2406.

Philippe H, Brinkmann H, Copley RR, et al. 2011. Acoelomorph flatworms are deuterostomes related to Xenoturbella. Nature. 470:255–258.

Philippe H et al. 2019. Mitigating Anticipated Effects of Systematic Errors Supports Sister-Group Relationship between Xenacoelomorpha and Ambulacraria. Curr. Biol. 29:1818–1826.e6.

Philippe H, Brinkmann H, Lavrov DV, et al. 2011. Resolving difficult phylogenetic questions: why more sequences are not enough. PLoS Biol. 9:e1000602.

R Core Team. 2013. R: A language and environment for statistical computing. http://r.meteo.uni.wroc.pl/web/packages/dplR/vignettes/intro-dplR.pdf.

Redmond AK, McLysaght A. 2021. Evidence for sponges as sister to all other animals from partitioned phylogenomics with mixture models and recoding. Nat. Commun. 12:1783.

Rokas A, Holland PW. 2000. Rare genomic changes as a tool for phylogenetics. Trends Ecol. Evol. 15:454–459.

Roure B, Rodriguez-Ezpeleta N, Philippe H. 2007. SCaFoS: a tool for selection, concatenation and fusion of sequences for phylogenomics. BMC Evol. Biol. 7 Suppl 1:S2.

Rouse GW, Wilson NG, Carvajal JI, Vrijenhoek RC. 2016. New deep-sea species of Xenoturbella and the position of Xenacoelomorpha. Nature. 530:94–97.

Ruiz-Trillo I, Paps J. 2016. Acoelomorpha: earliest branching bilaterians or deuterostomes? Org. Divers. Evol. https://link.springer.com/article/10.1007/s13127-015-0239-1.

Ruiz-Trillo I, Riutort M, Fourcade HM, Baguñà J, Boore JL. 2004. Mitochondrial genome data support the basal position of Acoelomorpha and the polyphyly of the Platyhelminthes. Mol. Phylogenet. Evol. 33:321–332.

Salichos L, Rokas A. 2011. Evaluating ortholog prediction algorithms in a yeast model clade. PLoS One. 6:e18755.

Schmidt HA, Strimmer K, Vingron M, von Haeseler A. 2002. TREE-PUZZLE: maximum likelihood phylogenetic analysis using quartets and parallel computing. Bioinformatics. 18:502–504.

Shen X-X, Hittinger CT, Rokas A. 2017. Contentious relationships in phylogenomic studies can be driven by a handful of genes. Nat Ecol Evol. 1:126.

Shen X-X, Salichos L, Rokas A. 2016. A Genome-Scale Investigation of How Sequence, Function, and Tree-Based Gene Properties Influence Phylogenetic Inference. Genome Biol. Evol. 8:2565–2580.

Shen X-X, Steenwyk JL, Rokas A. 2021. Dissecting incongruence between concatenation-and quartet-based approaches in phylogenomic data. Syst. Biol. doi: 10.1093/sysbio/syab011.

Shimodaira H. 2002. An approximately unbiased test of phylogenetic tree selection. Syst. Biol. 51:492–508.

Siu-Ting K et al. 2019. Inadvertent Paralog Inclusion Drives Artifactual Topologies and Timetree Estimates in Phylogenomics. Mol. Biol. Evol. 36:1344–1356.

Smith ML, Hahn MW. 2021. The frequency and topology of pseudoorthologs. bioRxiv. 2021.02.17.431499. doi: 10.1101/2021.02.17.431499.

Smith SA, Walker-Hale N, Walker JF, Brown JW. 2020. Phylogenetic Conflicts, Combinability, and Deep Phylogenomics in Plants. Syst. Biol. 69:579–592.

Spillane JL, LaPolice TM, MacManes MD, Plachetzki DC. 2021. Signal, bias, and the role of transcriptome assembly quality in phylogenomic inference. BMC Ecol Evol. 21:43.

Steenwyk JL et al. 2021. PhyKIT: a broadly applicable UNIX shell toolkit for processing and analyzing phylogenomic data. Bioinformatics. doi: 10.1093/bioinformatics/btab096.

Steenwyk JL, Buida TJ 3rd, Li Y, Shen X-X, Rokas A. 2020. ClipKIT: A multiple sequence alignment trimming software for accurate phylogenomic inference. PLoS Biol. 18:e3001007.

Struck TH. 2014. TreSpEx-Detection of Misleading Signal in Phylogenetic Reconstructions Based on Tree Information. Evol. Bioinform. Online. 10:51–67.

Thompson JD, Plewniak F, Ripp R, Thierry JC, Poch O. 2001. Towards a reliable objective function for multiple sequence alignments. J. Mol. Biol. 314:937–951.

Walker JF, Shen X-X, Rokas A, Smith SA, Moyroud E. 2020. Disentangling biological and analytical factors that give rise to outlier genes in phylogenomic matrices. doi: 10.1101/2020.04.20.049999.

Wilberg EW. 2015. What’s in an Outgroup? The Impact of Outgroup Choice on the Phylogenetic Position of Thalattosuchia (Crocodylomorpha) and the Origin of Crocodyliformes. Syst. Biol. 64:621–637.

Wilkinson M, McInerney JO, Hirt RP, Foster PG, Embley TM. 2007. Of clades and clans: terms for phylogenetic relationships in unrooted trees. Trends Ecol. Evol. 22:114–115.

Yu G, Smith DK, Zhu H, Guan Y, Lam TT-Y. 2017. ggtree: an r package for visualization and annotation of phylogenetic trees with their covariates and other associated data. Methods Ecol. Evol. 8:28–36.

Zwickl DJ, Hillis DM. 2002. Increased taxon sampling greatly reduces phylogenetic error. Syst. Biol. 51:588–598.

